# Long-term impact of early life stress on serotonin connectivity

**DOI:** 10.1101/2023.08.01.551573

**Authors:** Raksha Ramkumar, Moriah Edge-Partington, Kabirat Adigun, Yi Ren, Dylan J Terstege, Nazmus S Khan, Nahid Rouhi, Naila F Jamani, Mio Tsutsui, Jonathan R Epp, Derya Sargin

**Author notes:** These authors contributed equally (Raksha Ramkumar, Moriah Edge-Partington). **Corresponding author:** Derya Sargin Address: University of Calgary 2500 University Ave Dr NW T2N 1N4 Calgary, AB, Canada.

## Abstract

Chronic childhood stress is a prominent risk factor for developing mood disorders, yet mechanisms underlying this association remain unclear. Serotonin (5-HT) plays a crucial role in neurodevelopment and vulnerability to mood disorders. Maintenance of optimal 5-HT levels during early postnatal development is critical for the maturation of brain circuits. Developmental stress can alter the serotonin system, leading to chronic behavioral deficits. Yet, our understanding of the long-term impact of early life stress (ELS) on serotonin connectivity remains incomplete. Using a mouse model of chronic developmental stress, we sought to determine how ELS impacts brain-wide serotonin activity and behavior in adulthood. We established that adult female and male mice exposed to ELS during the first postnatal week show heightened anxiety-like behavior. Using *in vivo* fiber photometry and c-fos dependent activity mapping, we found that ELS enhances susceptibility to acute stress by disrupting the brain-wide functional connectivity of the raphe nucleus and the activity of dorsal raphe serotonin neuron population, in conjunction with a profound increase in the orbitofrontal cortex (OFC) activity. We further identified that 5-HT release in the medial OFC during environmental challenge is disrupted in mice exposed to ELS. Optogenetic stimulation of 5-HT terminals in the mOFC elicited an anxiolytic effect in ELS mice in a sex-dependent manner. Our findings hold significant insight into the mechanisms underlying long-term brain connectivity deficits induced by ELS, with potential implications for developing targeted stimulation-based treatments for affective disorders that arise from early life adversities.

## Introduction

Exposure to early life stress (ELS) is associated with changes in brain function, emotional behavior, and can contribute to the development of psychiatric disorders in adulthood^1–3^. Research over the past three decades has established the long-lasting effects of ELS on the risk and course of mood and anxiety disorders^4^. Although genetics can influence vulnerability to these disorders, environmental factors play a key role in contributing to worsened psychological and behavioral outcomes^5^. Early traumatic experiences including social deprivation, neglect, and abuse correlate with aberrant hypothalamic pituitary adrenal axis reactivity^6^, altered brain activity^7, 8^ and impaired social and emotional behaviors^9^, with an overall increased risk for lifetime mood disorders^10–12^.

In mammals, there are critical periods in development where exposure to stress can lead to enduring maladaptive psychological effects. After birth, the brain continues to undergo significant developmental processes such as neuronal growth, synaptic stabilization, and synaptic pruning^13^. The developing brain is highly plastic to allow for organization and adaptation to the environment, which consequently creates a vulnerability to external influences^14, 15^. Traumatic experience or chronic stress during this critical developmental period can impact brain function, behavior, and induce vulnerability to future stress. In particular, brain regions with extended postnatal development including the higher-order cortical structures, such as the dorsolateral prefrontal and orbitofrontal cortices are suggested to have increased vulnerability to the negative effects of ELS^16, 17^.

Serotonin (5-hydroxytryptamine; 5-HT) is a neurotransmitter that is widely distributed across the central nervous system (CNS) and is critical for social and emotional regulation. During early development, serotonin regulates cell survival, growth and differentiation, and is important for the maturation of neural circuits^18, 19^. Peak serotonin levels occur during the first two years in humans^20^ and the first postnatal week in rodents^21^. Manipulations that alter the serotonin system during these periods, such as stress, physical abuse, and lack of maternal care, are associated with chronic behavioral deficits in rodents and primates^15^. Maternal absence during development lowers serotonin release^22, 23^, disrupts serotonin receptor levels and function^24, 25^ resulting in mood deficits in adulthood^26–28^. Disruption of serotonin signaling via the blockade of 5-HT transporters or manipulation of 5-HT levels results in behavioral deficits during adulthood^29–31^. Some of these impairments can be rescued by increasing serotonin levels in the affected brain regions^27^, providing evidence of the role of 5-HT in the pathophysiology and treatment of these conditions.

5-HT neurons originate from the nine distinct nuclei clustered in the brainstem and project widely throughout the brain providing widespread modulation of the activity of many neural networks^32, 33^. The dorsal raphe nucleus (DRN) contains the majority of 5-HT-producing neurons which are highly heterogenous in terms of physiological function and gene expression profiles^32, 34, 35^. DRN 5-HT neurons densely innervate cortical and subcortical regions of the brain, regulating a wide range of physiological and behavioral processes^36, 37^. Emerging evidence implicates DRN 5-HT neurons to modulate emotionally salient information via diverse subcircuits. There is considerable overlap in the response of DRN 5-HT neurons to differentially salient stimuli however, distinct projection subpopulations can show functional bias based on their downstream connectivity. While a mixed population of DRN 5-HT neurons are activated by reward^35, 38–40^, those projecting to cortical and subcortical regions encode opposing responses to aversive stimuli^38^. Despite the involvement of each projection in emotional behavior, the proportion of DRN 5-HT neurons projecting to reward- or anxiety-modulating regions is correlated with the specific function of each pathway, leading to a projection-based average response to stimuli associated with positive or negative valence.

Overall, DRN 5-HT neurons play a critical role in environment-specific behavioral regulation when faced with positive stimuli or threat^41^. Early postnatal stress can disrupt the maturation and activity of the 5-HT circuits, leading to persistent negative effects on emotional behavior and threat response^42^. It is imperative to develop targeted treatments to overcome ELS-induced behavioral dysregulation. Yet, we do not fully understand how distinct 5-HT projections involved in emotionally salient behavior are impacted by ELS. Here, we first established a modified chronic ELS model based on the limited bedding and nesting (LBN) paradigm^43^ in female and male mice. When ELS offspring reached adulthood, we parsed out the threat-induced alterations in raphe nucleus functional connectivity, 5-HT neuron activity and consequent 5-HT release in mice exposed to ELS, compared with controls. We found that ELS during postnatal day 2-10 (PND2-10) disrupts functional connectivity of the raphe nucleus and perturbs the response of 5-HT neurons to threat, imposed by footshock. This is accompanied by an increased activity in the orbitofrontal cortex (OFC) but not central amygdala (CeA), two regions that receive dense 5-HT innervation and encode emotionally salient stimuli. Further analysis of the 5-HT signaling in the medial orbitofrontal cortex (mOFC) revealed blunted 5-HT release in the face of challenge in mice exposed to ELS, and a disruption of 5-HT signaling in the mOFC pyramidal neurons. Optogenetic stimulation of 5-HT projections in the mOFC improved anxiety-like behavior observed in male ELS mice, suggesting a sex-dependent anxiolytic effect. Ultimately, these findings have important implications for stimulation-based treatment approaches.

## Materials and Methods

Refer to Supplemental Methods for the extended details.

### Animals

Female and male Fos[2A-iCreER] (TRAP2) (JAX 030323) mice crossed with Ai14 reporter line (JAX 007914), and ePet1-cre (JAX 012712) mice crossed with Ai148 reporter line (JAX 030328) on C57Bl6 background were used for all experiments.

### Limited Bedding and Nesting

LBN was performed as previously described^43^ on PND2-10 with modifications in the protocol to enhance pup survival. Maternal behavior was recorded during light and dark cycles.

### Network connectivity and FASTMAP analysis

For brain-wide functional connectivity analyses, c-fos immunoreactive density was assessed across 60 neuroanatomical regions (see Table S1) using NeuroInfo software (MBF Bioscience). Regional c-fos immunoreactive density was cross-correlated across all mice within each group to generate correlation matrices of regional coactivation. For subsequent targeted c-fos immunoreactive density analyses, FASTMAP was used for the bilateral registration of the OFC and CeA.

### Stereotaxic Viral Delivery and Fiber Implantation

For fiber photometry experiments, 400 nl AAV2/9-CAG-iSeroSnFR-NLG (Canadian Neurophotonics Platform) was infused into the mOFC (AP: 2.34, ML: 0.3, DV: 2.5 mm from Bregma), followed by optic fiber implantation. For ePet1::Ai148 mice, the optic fiber was implanted above the DRN (30° angle, AP: -6.27, ML: 0, DV: -4.04 from Bregma). For optogenetic experiments, 800 nl AAV2/9-EF1a-DIO-mCherry or AAV2/9-EF1a-DIO-hChR2(H134R)-mCherry was infused into the DRN. Optic cannulae were implanted above the mOFC bilaterally (21° angle, AP: 2.34, ML: ±1.25, DV: -2.6 from Bregma).

### *In vivo* Fiber Photometry

Ca^2+^ signal of the 5-HT neuron population was recorded in ePet1::Ai148 mice to measure 5-HT neuron activity in response to footshocks (0.5 mA, 2 sec, 10X). Serotonin release in mOFC was recorded in mice infused with the iSeroSnFR 5-HT sensor during open field and tail suspension tests. Data were extracted and analyzed using a custom-written script in MATLAB (Mathworks).

### Statistical Analysis

Statistical analysis was performed using GraphPad Prism 9 software. Data were shown as means ± standard error of the mean (SEM). Data were analyzed using two-way ANOVA or unpaired t-test. Following significant interactions in two-way ANOVA, post-hoc analysis was conducted using Tukey’s multiple comparisons tests.

## Results

### LBN differentially affects dam behavior in light and dark cycles

To model early life stress (ELS), we used a modified limited bedding and bedding nesting (LBN) paradigm (**Figure 1A**). On postnatal day (PND) 2, dams and pups were placed into LBN cages until PND 10. Control dams and pups were moved to a cage that was otherwise identical but contained regular bedding and nesting material. The time dams spent with pups was analyzed using 30 min recordings during the light (day; 8:00-9:30 am) and dark (night; 9:00-10:30 pm) cycles separately on PND 3, 6 and 9. Control and LBN dams spent comparable time with pups during the day (two-way ANOVA; effect of group: F(1, 48) = 2.51, *p* = 0.12, effect of time: F(2, 48) = 0.86, *p* = 0.43) **(Figure 1B**). During night time, control dams spent significantly less time with pups compared with LBN dams, predominantly on PND 6 and 9 (two-way ANOVA; effect of group: F(1, 48) = 7.79, *p* = 0.007, effect of time: F(2, 48) = 2.84, *p* = 0.07) (**Figure 1C**). Although the total time spent with pups did not differ between control and LBN dams during the light cycle, dams exposed to LBN exhibited higher number of exits from the nests compared with control dams (two-way ANOVA; effect of group: F(1, 48) = 10.02, *p* = 0.003, effect of time: F(2, 48) = 0.35, *p* = 0.71) (**Figure 1D**). While the LBN dams spent significantly greater time with their pups during the dark cycle, the amount of nest exits/entries were significantly higher when compared with control dams (two-way ANOVA; effect of group: F(1, 48) = 19.95, *p* < 0.001, effect of time: F(2, 48) = 1.74, *p* = 0.19) (**Figure 1E**). During the time away from their pups, LBN dams were engaged with drinking and eating, similar to the control dams but also engaged in frequent bouts of tail chasing, a behavior that was not observed among control dams. Overall, dams exposed to LBN showed the most striking differences in maternal care during the dark cycle, with a significantly higher time spent with pups accompanied by greater frequency of nest entries and exits. This change in behavior may indicate attempted compensatory care on the part of the LBN dams to overcome the impoverished housing conditions.

**Figure 1.**
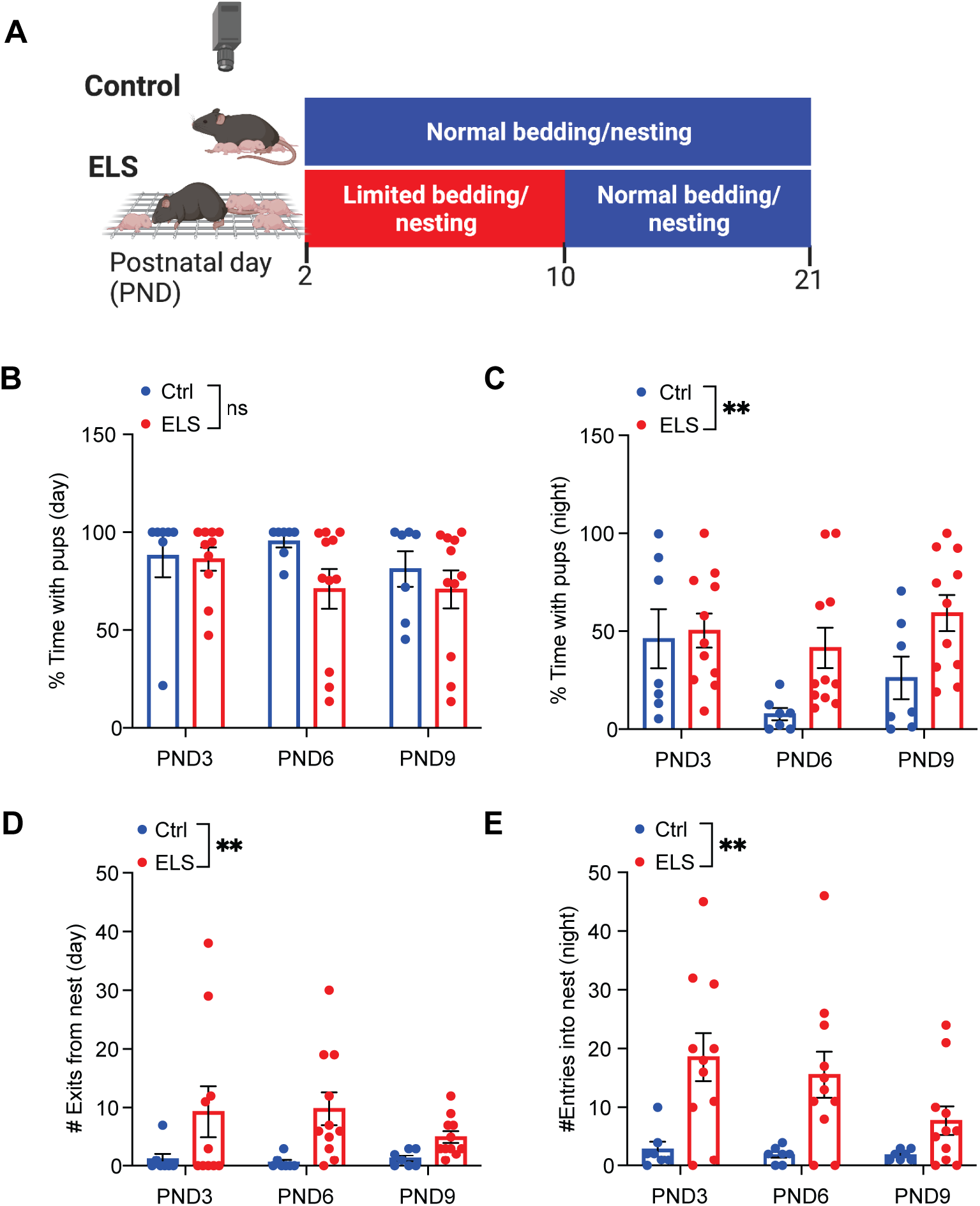
Maternal behavior during light and dark cycle. **A)** Experimental paradigm showing the implementation of control and limited bedding/nesting (LBN) conditions. Dams and pups were exposed to LBN on PND2-10. **B)** The time LBN dams (n = 11) spent with pups on PND3, 6 and 9 during the light cycle was comparable to control (n = 7) dams (two-way ANOVA; effect of group: F(1, 48) = 2.51, *p* = 0.12, effect of time: F(2, 48) = 0.86, *p* = 0.43). **C)** LBN dams spent significantly greater time with pups across PND3, 6 and 9 during the dark cycle, compared to control dams (two-way ANOVA; effect of group: F(1, 48) = 7.79, \*\**p* = 0.007, effect of time: F(2, 48) = 2.84, *p* = 0.07). **D)** LBN dams exhibited significantly higher frequency of next exits on PND3, 6 and 9 during the light cycle, compared to control dams (two-way ANOVA; effect of group: F(1, 48) = 10.02, \*\**p* = 0.003, effect of time: F(2, 48) = 0.35, *p* = 0.71). **E)** During the dark cycle, LBN dams exhibited significantly higher frequency of nest entries, compared to control dams (two-way ANOVA; effect of group: F(1, 48) = 19.95, \*\**p* < 0.01, effect of time: F(2, 48) = 1.74, *p* = 0.19). \*\**p*<0.01, ns; non-significant. Data represent mean ± SEM.

### Female and male offspring exposed to LBN show anxiety-like behavior in adulthood

To determine the long-term impacts on the behavior of offspring reared under ELS conditions, we performed open field, 3-chamber social interaction and tail suspension tests when pups reached adulthood (> PND 60) (**Figure 2A**). Female and male mice exposed to ELS spent a significantly greater amount of time in the outer zone of the open field (**Figure 2B**; two-way ANOVA; effect of group: F(1, 34) = 19.69, *p* < 0.001, effect of sex: F(1, 34) = 1.48, *p* = 0.23) while they spent significantly less time in the intermediate (**Figure 2C**; two-way ANOVA; effect of group: F(1, 34) = 16.84, *p* < 0.001, effect of sex: F(1, 34) = 1.77, *p* = 0.19) and inner (**Figure 2D**; two-way ANOVA; effect of group: F(1, 34) = 13.52, *p* < 0.001, effect of sex: F(1, 34) = 0.006, *p* = 0.94) zones, indicating greater anxiety-like behavior, compared with control female and male mice. The overall distance traveled in the open field also differed between control and ELS mice, with significant sex-dependent effects (**Figure S1A**; two-way ANOVA; effect of group: F(1, 34) = 4.31, *p* = 0.04, effect of sex: F(1, 34) = 16.4, *p* < 0.001).

**Figure 2.**
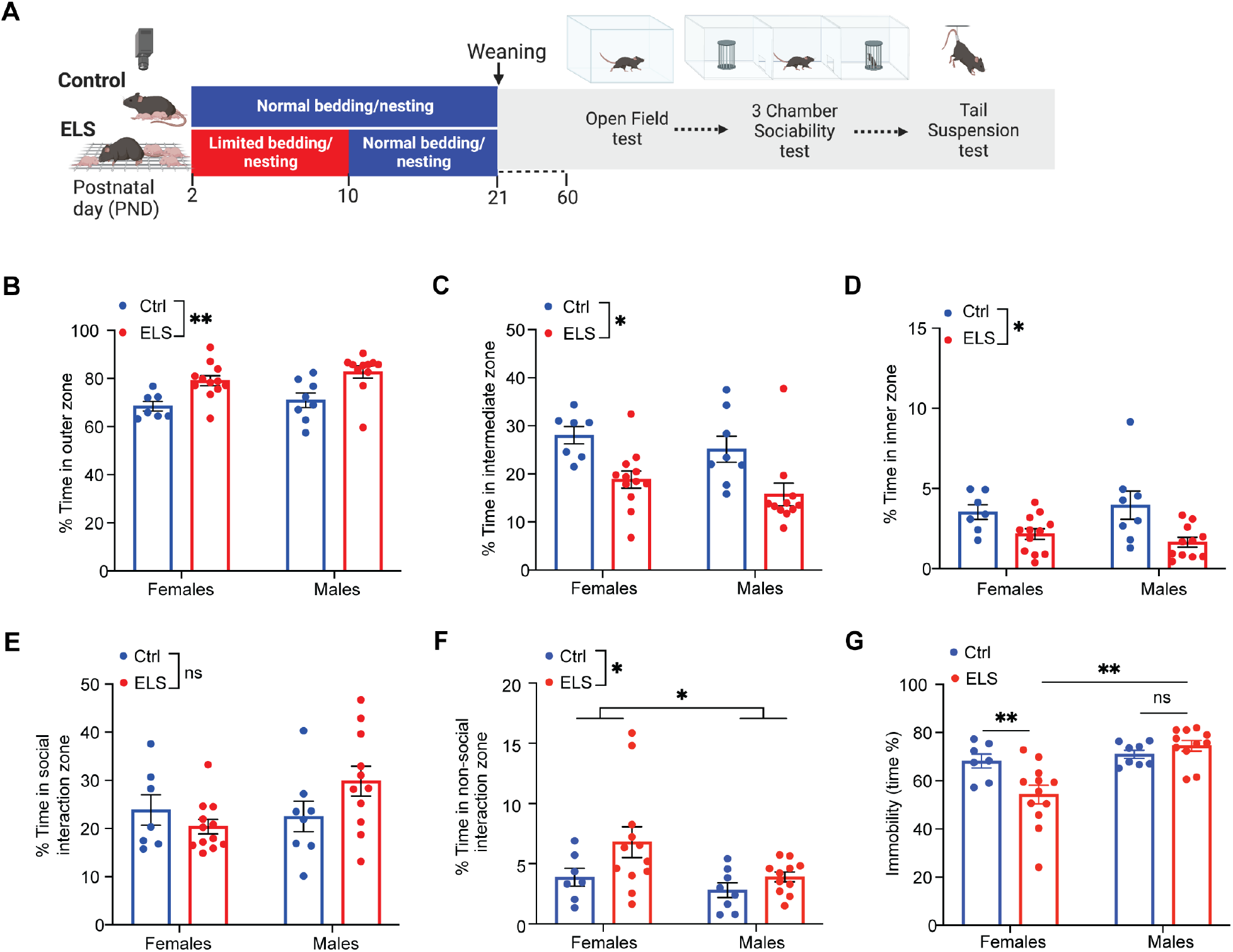
Long-term impact of ELS on offspring behavior. **A)** Experimental paradigm showing behavioral tests performed after offspring reared under control and LBN conditions reached adulthood (> PND60). Female (n = 12) and male ELS (n = 11) mice spent significantly greater amount of time in the outer zone (two-way ANOVA; effect of group: F(1, 34) = 19.69, \*\**p* < 0.001, effect of sex: F(1, 34) = 1.48, *p* = 0.23) **(B)**, and significantly less time in the intermediate (two-way ANOVA; effect of group: F(1, 34) = 16.84, \*\**p* < 0.001, effect of sex: F(1, 34) = 1.77, *p* = 0.19) **(C)** and inner (two-way ANOVA; effect of group: F(1, 34) = 13.52, *p* < 0.001, effect of sex: F(1, 34) = 0.006, *p* = 0.94) **(D)** zones of the open field, indicating a greater anxiety-like behavior, compared with female (n = 7) and male (n = 8) controls. **E)** Time spent interacting with a stranger mouse (social interaction zone) was comparable between female and male control and ELS mice (two-way ANOVA; effect of group: F(1, 34) = 0.49, *p* = 0.49, effect of sex: F(1, 34) = 2.15, *p* = 0.15). **F)** Female and male ELS mice spent significantly greater time interacting with an empty cup (non-social interaction zone) compared to control mice (two-way ANOVA; effect of group: F(1, 34) = 4.47, \**p* = 0.04, effect of sex: F(1, 34) = 4.35, \**p* = 0.04). **G)** Female and male ELS mice show frequently greater passive coping behavior (immobility) during tail suspension test, compare to controls (two-way ANOVA; effect of group: F(1, 34) = 17.4, \*\**p* < 0.01, effect of sex: F(1, 34) = 7.57, \*\**p* = 0.009). \**p*<0.05, \*\**p*<0.01, ns; non-significant. Data represent mean ± SEM.

Next, we performed 3-chamber social interaction test to assess the effect of ELS on sociability. The time spent interacting with a sex- and age-matched stranger conspecific was comparable between ELS and control mice (two-way ANOVA; effect of group: F(1, 34) = 0.49, *p* = 0.49, effect of sex: F(1, 34) = 2.15, *p* = 0.15) (**Figure 2E**). However, female and male ELS mice spent a significantly greater time in the non-social interaction zone, investigating the empty cup compared with controls (two-way ANOVA; effect of group: F(1, 34) = 4.47, *p* = 0.04, effect of sex: F(1, 34) = 4.35, *p* = 0.04) (**Figure 2F**). The sociability index showed a significant sex difference between female and male mice, but female and male ELS groups showed comparable sociability to controls (two-way ANOVA; effect of group: F(1, 34) = 3.21, *p* = 0.08, effect of sex: F(1, 34) = 6.11, *p* = 0.019) (**Figure S1B**). Female ELS mice showed a tendency to spend less time in the social chamber (two-way ANOVA; group X sex interaction: F(1, 34) = 5.62, *p* = 0.024, Tukey’s multiple comparisons test: Female ctrl vs ELS *p* = 0.067; female ELS vs male ELS *p* = 0.06, all other *p*’s > 0.1) (**Figure S1C**) and spent significantly greater time in the non-social chamber, compared with controls (two-way ANOVA; group X sex interaction: F(1, 34) = 8.18, *p* = 0.007, Tukey’s multiple comparisons test: Female ctrl vs ELS *p* = 0.04; female ELS vs male ELS *p* = 0.007, all other *p*’s > 0.1) (**Figure S1D**).

In order to investigate the effect of a stressful environment on coping behaviors in both control and ELS mice, we conducted a tail suspension test (TST), which acts as an inescapable stressor in a high threat environment^41^. We quantified the time mice were engaged in active (struggling behavior) and passive (immobility) coping behaviors^44^ over the entire duration of the test. Female ELS mice spent significantly greater time engaged in active coping behavior indicated by increased mobility duration, compared with control females and ELS males. We did not find a difference in stress coping strategies in TST between control and ELS males (two-way ANOVA; group X sex interaction: F(1, 34) = 7.78, *p* = 0.009, Tukey’s multiple comparisons test: Female ctrl vs ELS *p* = 0.02; female ELS vs male ELS *p* < 0.001, male ctrl vs ELS *p* = 0.84) (**Figure 2G**).

Together, these data suggested that female and male offspring exposed to ELS show striking increases in anxiety-like behavior. Sociability is reduced only in female ELS mice while both male control and ELS mice spent comparable time interacting with strangers. Female ELS mice additionally show greater inescapable stress-induced active coping behavior during adulthood.

### ELS disrupts raphe nucleus functional connectivity

ELS has been associated with disrupted brain connectivity and increased emotional and social deficits later in life^7, 45, 46^. The organization of behavior is highly complex and most functions tend to be distributed throughout numerous regions of the brain. In order to understand ELS-induced functional network deficits, we sought to determine changes in the patterns of functional connectivity in response to threat in control and ELS mice. We used FosTRAP2::tdTomato mice in which cells that are active during a discrete temporal window can be permanently tagged with the fluorescent reporter tdTomato^47^. FosTRAP2::tdTomato mice were administered ten 0.5 mA foot shocks prior to intraperitoneal (i.p.) injection of 4-hydroxytamoxifen. Mice were perfused three days later and entire brains were cryosectioned, counterstained and imaged. c-Fos-immunoreactive cells were segmented using supervised machine learning and images were registered to the Allen Mouse Brain Reference Atlas. Regional c-fos-immunoreactive densities were correlated across mice within each group, to generate correlated activity matrices (**Figure 3A**). We then examined functional connectivity based on the regional c-fos immunoreactivity within 60 brain regions. Because the raphe nucleus is strongly modulated in response to stress and important for emotional regulation, we used this region as a seed to assess its functional connectivity in both female and male control and ELS mice exposed to footshock stress (**Figure 3B**). The number of brain regions showing anti-correlated activity with the raphe nucleus was greater in ELS mice, compared to controls, with the most striking differences present in the male ELS group (**Figure 3C, D, Table S1**). This suggested that ELS leads to a large increase in anti-correlated functional connectivity of the raphe nucleus signifying a loss of coordinated activity between the raphe nucleus and many other brain regions. We plotted the distribution of the Pearson’s correlation coefficients (*R* values) for all the possible correlations in each network. The comparison of the distribution of *R* values revealed significant differences in the distribution between male control and ELS mice (K-S test, *p* < 0.0001), female control and ELS mice (K-S test, *p* = 0.0025) as well as between male ELS and female ELS mice (K-S test, *p* < 0.0001). Importantly we did not see a difference in the distribution of *R* values between male and female control groups (K-S test, *p* = 0.18) (**Figure 3E**).

**Figure 3.**
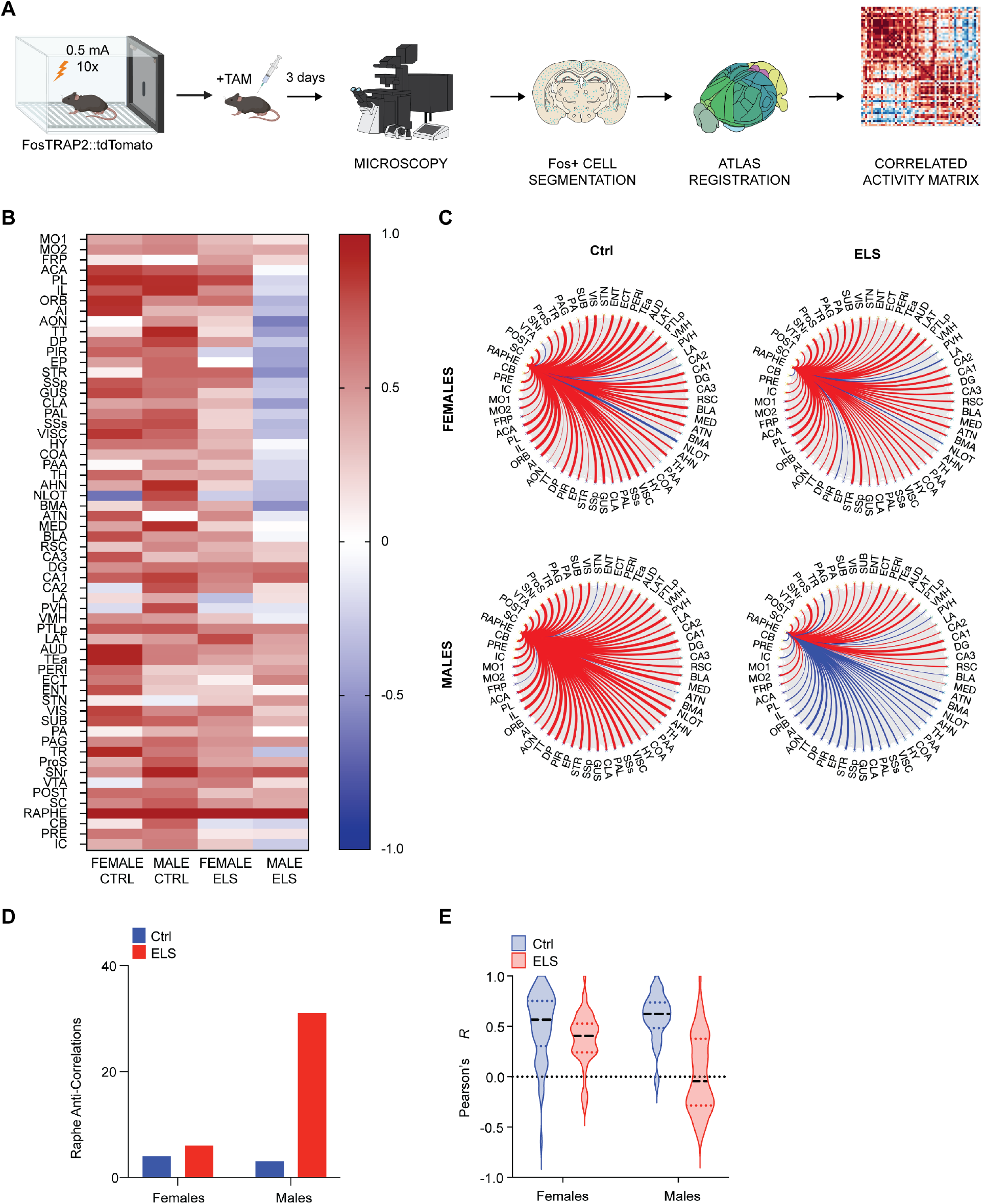
ELS-induced disruption in raphe functional connectivity. **A)** Pipeline for assessing functional connectivity underlying acute footshock stress in female control (n = 7), male control (n = 7), female ELS (n = 11) and male ELS (n = 10) mice. **B)** Functional connectivity of the raphe nucleus based on the regional c-fos activity was assessed by isolating the column in the correlated activity matrix corresponding to this region for female and male control and ELS mice. **C)** ELS led to an increase in anti-correlated connectivity in the raphe nucleus, which was more prominent in males. Blue lines depict anti-correlations, red lines show positive correlations. Line weight is indicative of the magnitude of the Pearson’s R value. **D)** Number of brain regions showing anti-correlated activity (negative correlations) with the raphe nucleus in female and male control and ELS mice. **E)** Distribution of Pearson correlation coefficients showed significantly altered correlation between control and ELS mice, with male ELS group showing the most striking difference from controls (Kolmogorov-Smirnov tests: Female Ctrl vs. Female ELS, *p* = 0.0025; Male Ctrl vs. Male ELS, *p* < 0.0001; Female Ctrl vs. Male Ctrl, *p* = 0.1813; Female ELS vs. Male ELS, *p* < 0.0001). Also see Table S1 for the list of names for abbreviated brain regions.

Taken together, the functional connectome of the raphe nucleus in ELS mice showed a higher number of anti-correlated functional connections, with males showing greater disruption in connectivity compared to females. These data suggest that offspring exposed to ELS show reduced synchronization between the raphe nucleus and other brain regions in adulthood.

### ELS disrupts serotonin neuron activity in response to an aversive stimulus

The raphe nucleus contains a heterogenous neural network composed predominantly of serotonin neurons, which play a crucial role in mediating threat adaptive behaviors^41^. Given the imperative role of serotonin during development, supported by our findings in ELS-induced disruption in raphe functional connectivity, we next determined the threat-induced differences in the activity of serotonin neuron population between control and ELS mice. For these experiments, we crossed the ePet1-Cre transgenic driver mouse line^48^ with the Ai148 reporter line to express the genetically encoded Ca^2+^ sensor GCaMP6f specifically in serotonin neurons (**Figure 4A, B**). We implanted the optical fiber above the dorsal raphe nucleus (DRN) and three weeks after surgery, we exposed control and ELS mice to acute footshock while recording the response of serotonin neuron population using fiber photometry. After a 3 min baseline recording, ten footshocks (0.5 mA, 2 sec duration each) were delivered with 30 second intervals (**Figure 4C**). The averaged Ca^2+^ signal from serotonin neuron population yielded a biphasic response to footshock with an initial increase, followed by a depression in activity in female and male control and ELS mice (**Figure 4D, E**). We analyzed and compared the footshock-induced increase (peak) and depression (trough) in the amplitude of the serotonin Ca^2+^ signal between each group. In females, two-way ANOVA revealed no significant interaction between group (control vs ELS) and time (baseline t=0 s; peak, t = 0-1 s; trough, t = 2-4 s) [F(2, 30) = 17.4, *p* = 0.55)]. In females, there was no significant main effect of group [F(1, 30) = 2.3, *p* = 0.14)] however, there was a significant main effect of time [F(2, 30) = 71.32, *p* < 0.001)], indicating footshock-induced changes in activity over time (**Figure 4F**). We found a significant main effect of group [F(1, 27) = 4.51, *p* = 0.04] and time [F(2, 27) = 32.48, *p* < 0.001] but no significant interaction between group and time in males (two way ANOVA; F(2, 27) = 1.7, *p* = 0.2) (**Figure 4G**). The quantification of footshock-induced maximum amplitude (peak) of the serotonin Ca^2+^ signal revealed no significant differences between the groups (two-way ANOVA; effect of group: F(1, 19) = 1.01, *p* = 0.33, effect of sex: F(1, 19) = 0.14, *p* = 0.72) (**Figure 2H**). The analysis of the magnitude of depression (trough) showed a significantly larger reduction in footshock-induced serotonin Ca^2+^ signal in female and male ELS mice, compared to controls (two-way ANOVA; effect of group: F(1, 19) = 12.14, *p* = 0.002, effect of sex: F(1, 19) = 2.11, *p* = 0.16) (**Figure 2I**). These data suggested that serotonin neurons show significantly reduced depression in activity in response to threat in ELS mice, compared with controls.

**Figure 4.**
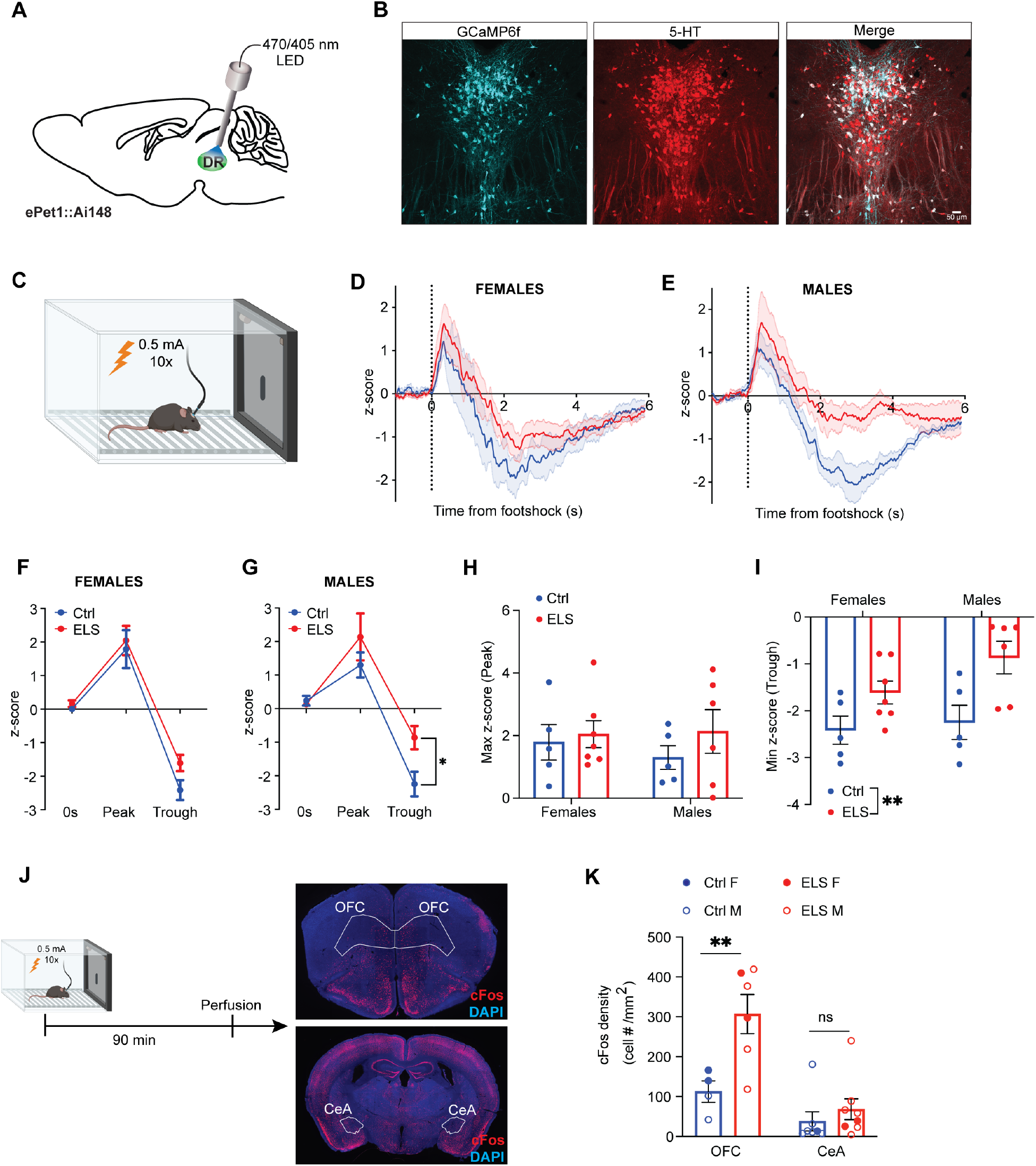
ELS alters footshock-induced activity of DR serotonin neurons and postsynaptic regions. **A)** Schematic showing implantation of an optical fiber in the dorsal raphe nucleus (DR) of ePet1::Ai148 mice, allowing *in vivo* Ca2+ imaging in serotonin neurons. **B)** Representative images showing GCaMP6f (cyan) and serotonin (red) co-localization in the DR of a ePet1::Ai148 mouse. **C)** Footshock paradigm (ten 0.5 mA footshocks, each 2 sec, presented with 30sec intervals). Averaged Ca^2+^ signal changes in serotonin neurons in female (ctrl: n = 5; ELS: n = 7) **(D)** and male (ctrl: n = 5; ELS: n = 6) mice **(E)** in response to acute footshock. The activity (z-score) of serotonin neuron population in response to footshock. The baseline at footshock onset (t=0s), the maximum response during t = 0-1 s (peak) and minimum response during t = 2-4 s (trough) were plotted for female **(F)** and male **(G)** control and ELS mice (two-way ANOVA; Females, effect of group: F(1, 30) = 2.3, *p* = 0.14), effect of time: F(2, 30) = 71.32, \*\**p* < 0.001; Males, effect of group: F(1, 27) = 4.51, \**p* = 0.04, effect of time: F(2, 27) = 32.48, \*\**p* < 0.001). **H)** Footshock-induced maximum serotonin neuron activity (peak) is comparable between female and male control and ELS mice (two-way ANOVA; effect of group: F(1, 19) = 1.01, *p* = 0.33, effect of sex: F(1, 19) = 0.14, *p* = 0.72). **I)** Footshock-induced minimum serotonin neuron activity (trough) is significantly reduced in female and male ELS mice, compared to controls (two-way ANOVA; effect of group: F(1, 19) = 12.14, \*\**p* = 0.002, effect of sex: F(1, 19) = 2.11, *p* = 0.16). **J)** Experimental paradigm showing representative images of c-fos staining in mice perfused 90 min after footshock. OFC, orbitofrontal cortex; CeA, central amygdala. **K)** ELS mice show increased activity in the OFC indicated by significantly greater number of c-fos positive cells induced by footshock, compared to controls. The activity in CeA is comparable between control and ELS mice (two-way ANOVA, group X brain region: F(1, 21) = 5.77, \**p* = 0.02; Tukey’s multiple comparisons test, OFC ctrl vs ELS: \*\**p* = 0.008, CeA ctrl vs ELS: *p* = 0.89). \**p*<0.05, \*\**p*<0.01, ns; non-significant. Data represent mean ± SEM.

The biphasic changes in serotonin neuron activity in response to footshock may be attributed to the heterogeneity of this population based on connectivity. Previous work has demonstrated that DRN serotonin neurons that project to the orbitofrontal cortex (OFC) and central amygdala (CeA) exhibit contrasting changes in activity in response to aversive stimuli^38^. Accordingly, serotonin neurons projecting to the CeA display a predominantly biphasic response to footshock, characterized by a large initial elevation in Ca^2+^ signal, followed by a small depression. In contrast, serotonin neurons projecting to the OFC predominantly exhibit a reduction in Ca^2+^ signal in response to footshock. In order to determine whether acute footshock modulates the activity of these projection regions differentially in control and ELS mice, we perfused a group of mice 90 min after footshock and immunolabeled the brains for c-fos (**Figure 4J**). We used the Flexible Atlas Segmentation Tool for Multi-Area Processing (FASTMAP) machine-learning pipeline^49^ for high-throughput quantification of c-fos density in the OFC and CeA from female and male control and ELS mice subjected to footshock. Two-way ANOVA revealed a significant interaction between group (ctrl, ELS) and brain region (OFC, CeA) [F(1, 21) = 5.77, *p* = 0.02]. Tukey’s multiple comparisons test showed that ELS mice have significantly more c-fos-immunoreactive cells in the OFC (Ctrl vs ELS, *p* = 0.008) while the c-fos density in the CeA was comparable between control and ELS mice (*p* = 0.89) (**Figure 4K**).

Together these data indicated that female and male mice subjected to ELS show reduced footshock-induced depression in serotonin Ca^2+^ signal, suggesting a disruption in serotonin response to acute stress. The analysis of c-fos immunoreactive density revealed that there are regional differences in footshock-induced activity following ELS. In the OFC, c-fos immunoreactivity is significantly more dense in ELS mice compared to controls, while activity in the CeA is similar between the two groups.

### ELS-induced disruption in serotonin release in avoidable and unavoidable threat environment

To determine whether activity dependent alterations in the orbitofrontal cortex of ELS mice could be mediated in part by stress-induced deficits in serotonin modulation, we investigated serotonin release during low- and high-threat environments^41^. We injected AAV2/9-CAG-iSeroSnFR-NLG^50^ expressing the serotonin sensor into the medial orbitofrontal cortex (mOFC) of control and ELS female and male C57Bl6 mice and implanted the optical fiber above the mOFC (**Figure 5A, B**). Four weeks after viral infusion, we performed fiber photometry while mice investigated an avoidable threat environment (open field). One week after the open field test, we performed fiber photometry while mice were subjected to an unavoidable threat (TST) (**Figure 5C**).

**Figure 5.**
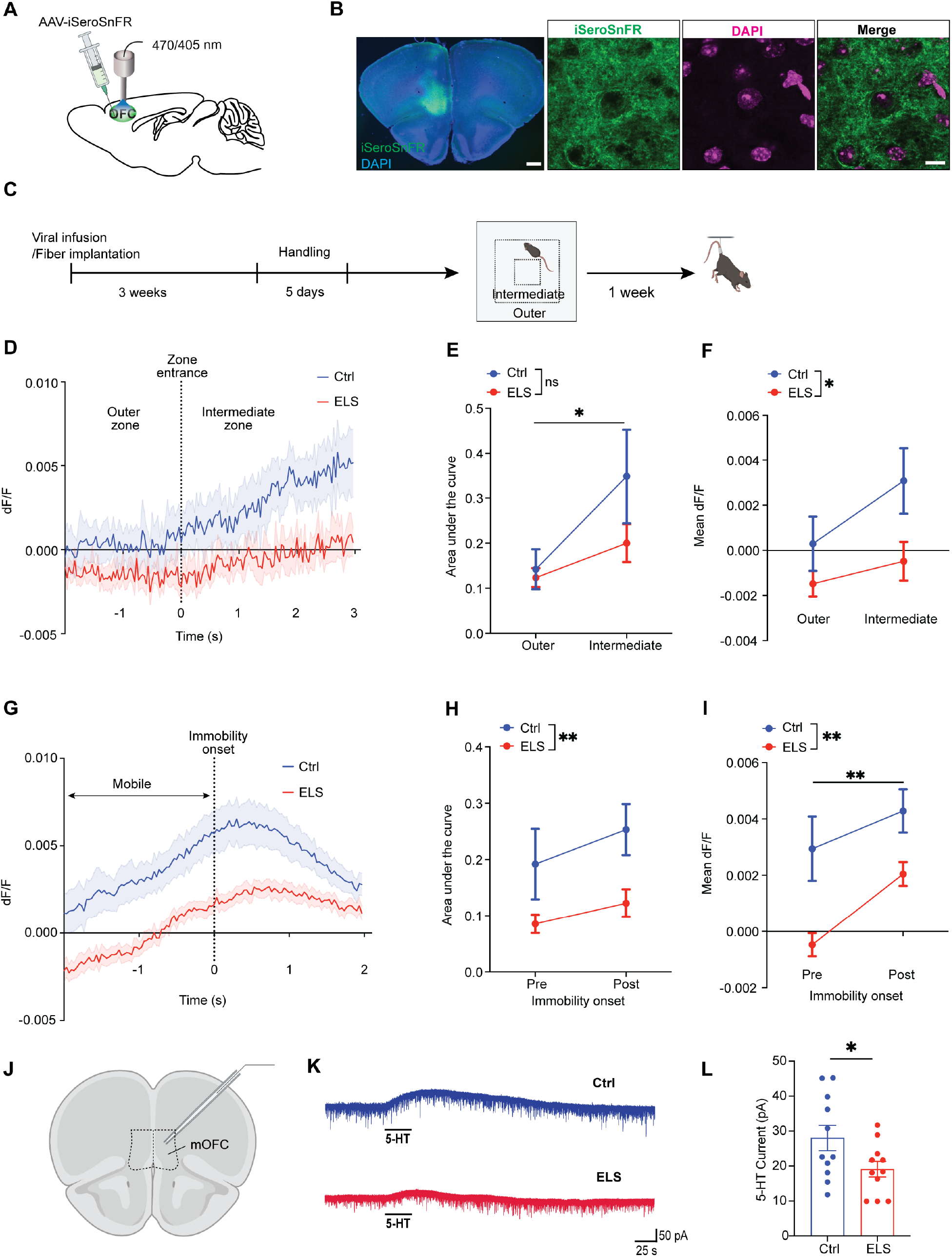
ELS leads to reduced serotonin release in the medial orbitofrontal cortex under aversive conditions. **A)** Schematic showing AAV2/9-CAG-iSeroSnFR-NLG infusion and fiber optic implantation in the medial orbitofrontal cortex (mOFC). **B)** Representative images showing iSeroSnFR (green) expression in the mOFC. Section is co-labeled with DAPI. **C)** Experimental scheme showing viral infusion and fiber implantation followed by handling prior to open field and tail suspension tests. **D)** Averaged iSeroSnFR signal 2 sec before and 3 sec after the mouse transitioned from the outer to the intermediate zone (ctrl: n = 7; ELS: n = 9). **E)** The area under the curve of the iSeroSnFR signal in the outer and intermediate zone. 5-HT is released in the mOFC after control and ELS mice entered the intermediate zone (two-way ANOVA; effect of group: F(1, 28) = 2.2, *p* = 0.15, effect of zone: F(1, 28) = 6.31, \**p* = 0.02). **F)** The mean amplitude of the iSeroSnFR signal (dF/F) in the outer and intermediate zone. ELS mice exhibit significantly lower serotonin release in the mOFC in the transition from outer to intermediate zone (two-way ANOVA; effect of group: F(1, 28) = 6.9, \*\**p* = 0.01, effect of zone: F(1, 28) = 3.48, *p* = 0.07). **G)** Averaged iSeroSnFR signal 2 sec before and after immobility onset in the tail suspension test (ctrl: n = 7; ELS: n = 9). **H)** The area under the curve of the iSeroSnFR signal before and after immobility onset in TST is significantly lower in ELS mice (two-way ANOVA; effect of group: F(1, 28) = 9.91, \*\**p* = 0.004; effect of mobility: F(1, 28) = 1.7, *p* = 0.2). **I)** The mean amplitude of the iSeroSnFR signal (dF/F) before and after immobility onset in TST is significantly lower in ELS mice (two-way ANOVA; effect of group: F(1, 28) = 16.82, \*\**p* < 0.001; effect of mobility: F(1, 28) = 7.9, \*\**p* = 0.009), suggesting reduced serotonin release. **J)** Slices comprising the mOFC were used for patch clamp recordings. **K)** Representative voltage-clamp recordings from mOFC neurons of a ctrl and an ELS mouse showing the outward inhibitory current in response to serotonin (5-HT) application. **L)** The amplitude of the outward inhibitory serotonin current is significantly lower in ELS mice, compared with controls (4 control male mice, n = 11 neurons; 3 ELS male mice, n = 11 neurons) (unpaired t-test, \**p* = 0.048). \**p*<0.05, \*\**p*<0.01, ns; non-significant. Data represent mean ± SEM.

Mice tend to perceive the inner zones of the open field as more aversive than the peripheral zones, likely due to the higher level of uncertainty. To investigate how serotonin release is modulated during transition from a lower to higher risk environment, we quantified the iSeroSnFR signal as mice moved from the outer zone to the intermediate zone (**Figure 5D**). The area under the curve of the iSeroSnFR signal in control and ELS mice was greater while mice were in the intermediate zone, compared with the outer zone (two-way ANOVA; effect of zone: F(1, 28) = 6.31, *p* = 0.02), suggesting greater serotonin release as mice transitioned to a more aversive environment (**Figure 5E, Figure S2A-C**). Although the area under the curve was not significantly different between the groups [F(1, 28) = 2.2, *p* = 0.15], the mean amplitude of the iSeroSnFR signal before and after entry into the intermediate zone revealed a significant reduction in mOFC serotonin release in ELS mice compared to control mice (two-way ANOVA; effect of group: F(1, 28) = 6.9, *p* = 0.01, effect of zone: F(1, 28) = 3.48, *p* = 0.07) (**Figure 5F, Figure S2A, B, D**).

TST exposes mice to an inescapable stressor, eliciting passive and active coping behaviors in response. Building on our observations of enhanced mobility in ELS mice during TST and the established link between chronic stress and altered coping strategies, we quantified the mOFC iSeroSnFR signal 2 sec before and after the onset of immobility in mice (**Figure 5G**). Compared to controls, ELS mice exhibited a significant decrease in both the area under the curve and mean amplitude of the iSeroSnFR signal before and after the onset of immobility in TST (AUC: two-way ANOVA; effect of group: F(1, 28) = 9.91, *p* = 0.004; mean dF/F: two-way ANOVA; effect of group: F(1, 28) = 16.82, *p* < 0.001) (**Figure 5H, I, Figure S2E-G**), with an overall increase in amplitude following immobility onset in both groups (two-way ANOVA; effect of mobility: F(1, 28) = 7.86, *p* = 0.009) (**Figure 5I, Figure S2E, F, H**). Our results indicate that serotonin release in the mOFC is greater during the transition to passive coping behavior in the TST, however this release is significantly lower in ELS mice compared to control, regardless of the coping strategy employed.

To investigate the physiological response of mOFC pyramidal neurons to serotonin, we next performed whole-cell patch-clamp electrophysiology in cortical slices obtained from control and ELS mice (**Figure 5J**). Voltage-clamp recordings revealed an inhibitory outward current in response to exogenous serotonin application in mOFC pyramidal neurons of control and ELS mice (**Fig 5K**). The amplitude of the outward serotonin current was significantly smaller in mOFC neurons of ELS mice compared to that of controls (unpaired t-test, *p* = 0.048). The passive membrane characteristics of mOFC neurons including the membrane capacitance (Cm), resting membrane potential (Vm) and input resistance (Rinput) did not differ between control and ELS mice (**Figure S3A-D**). The spike threshold was significantly more depolarized in mOFC neurons of ELS mice (unpaired t-test, *p* = 0.046), without changes in spike amplitude or intrinsic excitability when compared with mOFC neurons of control mice (**Figure S3E-H**).

Taken together, our data showed that ELS is associated with a significant reduction in mOFC serotonin release during high risk and high threat conditions. Additionally, we observed a reduced inhibitory response to serotonin in mOFC pyramidal neurons of ELS mice, indicating a disruption in the long-term serotonin-induced modulation of mOFC activity caused by ELS.

### Optogenetic stimulation of serotonin terminals in mOFC elicits an anxiolytic effect in male ELS mice

The orbitofrontal cortex is integral to encoding emotionally salient stimuli^51^. Altered activity of the OFC has been associated with anxiety disorders^52^, especially during exposure to anxiety-inducing imagery in humans^53, 54^, while OFC lesions lead to dysregulated threat response and anxiety, as shown in primates^55^. Successful antidepressant treatment targeting 5-HT can overcome deficits in OFC activity^56, 57^. However, the role of the DR^5-HT^ è mOFC pathway in emotional behavior remains to be determined. To investigate whether stimulation of 5-HT terminals in mOFC in adult female and male mice exposed to ELS is sufficient to deficits in emotional behavior, we infused a Cre-dependent AAV encoding channelrhodopsin-2 fused to mCherry (hChR2-mCherry) or mCherry only (control) into the DR of ePet1-Cre transgenic mice and implanted the optical fibers bilaterally over the mOFC to selectively illuminate this region (**Figure 6A**). This led to robust expression of the virus in the DR 5-HT neurons and their processes in the mOFC (**Figure 6B, C**). Using slice electrophysiology, we verified that the activation of 5-HT neuron terminals in the mOFC with blue light resulted in the suppression of action potentials in pyramidal neurons, which was abolished by application of the 5-HT1A receptor antagonist WAY-100635 (**Figure 6D**). 5 weeks after surgeries, we performed open field test followed by TST in male and female ELS mice infused with mCherry or hChR2-mCherry (**Figure 6E**) while stimulating the 5-HT neuron projections in the mOFC using blue light (470 nm, 20 Hz, 10 ms, 3 min epochs). In the open field, optogenetic stimulation of 5-HT projections in the mOFC increased the time spent in the outer zone, and decreased the time in the central zone in male ELS mice, suggesting an anxiolytic effect (two-way ANOVA; outer zone, effect of group: F(1, 40) = 16.59, *p* = 0.0002, effect of time: F(4, 40) = 0.62, *p* = 0.65; central zone, effect of group: F(1, 40) = 16.59, *p* = 0.0002, effect of time: F(4, 40) = 0.59, *p* = 0.67) (**Figure 6F, G**). In contrast, in female ELS mice, we did not obtain a difference between ChR2 and mCherry groups in the open field test upon optogenetic stimulation of the mOFC 5-HT terminals (two-way ANOVA; outer zone, effect of group: F(1, 40) = 0.77, *p* = 0.39, effect of time: F(4, 40) = 0.47, *p* = 0.76) (**Figure 6H, I**). We also did not obtain any changes in locomotor activity upon blue light stimulation in female and male ChR2 and mCherry mice in the open field test (two-way ANOVA; change in distance traveled light on – light off; effect of group: F(1, 16) = 0.03, *p* = 0.87, effect of sex: F(1, 16) = 1.76, *p* = 0.2).

**Figure 6.**
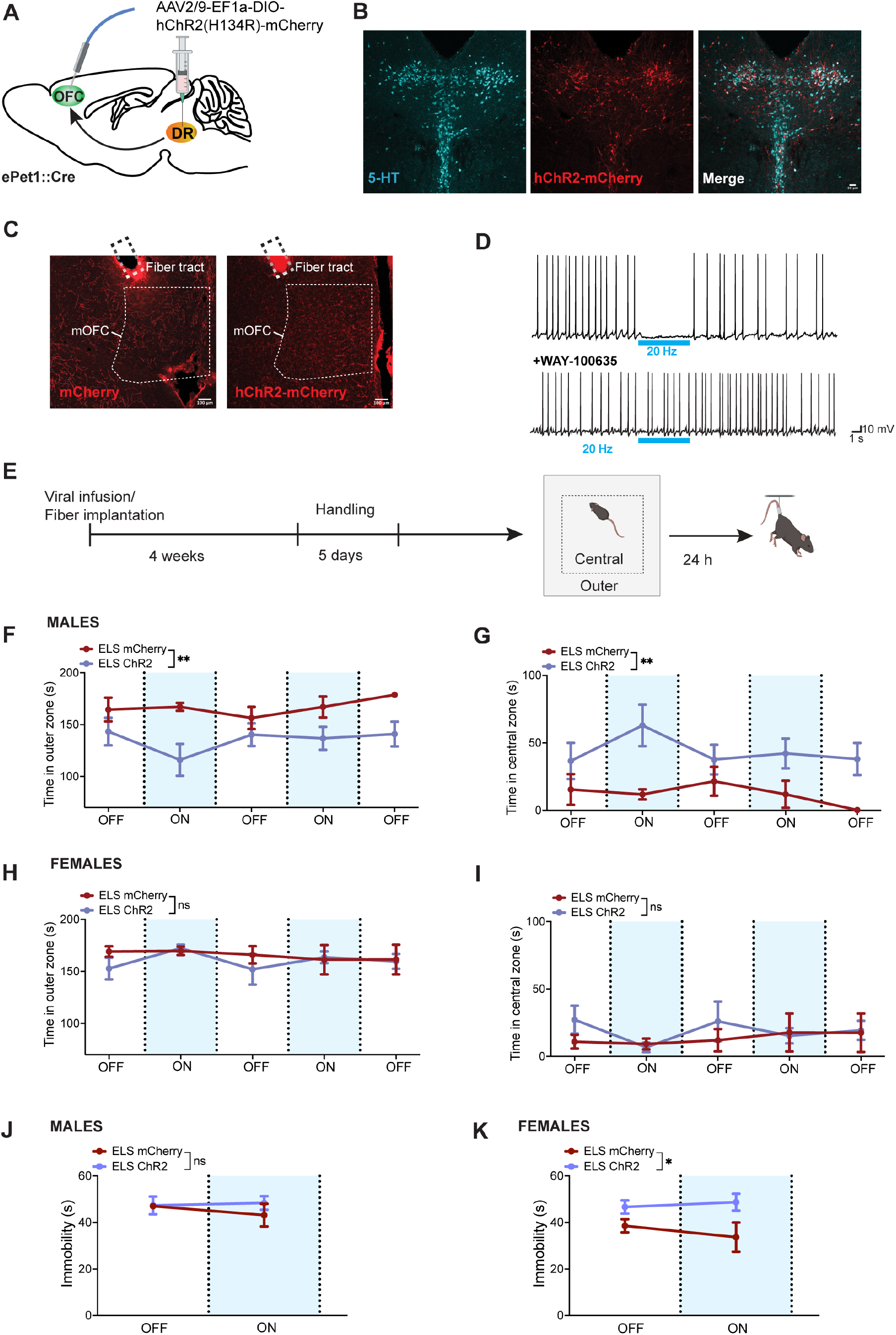
Optogenetic stimulation of 5-HT terminals in mOFC improves anxiety-like behavior in male ELS mice. **A)** Schematic showing AAV2/9-EF1a-DIO-hChR2(H134R)-mCherry infusion in DR and fiber optic implantation in mOFC. **B)** Representative images showing 5-HT (cyan) and hChR2-mCherry (red) expression in DRN. Scale bar, 50 µm. **C)** Representative images showing mCherry-labeled 5-HT neuron processes in the mOFC from mice expressing mCherry or hChR2-mCherry. Scale bars, 100 µm. **D)** Representative current clamp traces from an mOFC pyramidal neuron showing inhibition of action potentials upon blue light (above; 470 nm, 20 Hz, 10 ms) stimulation and the absence of light-induced inhibition after application of the 5-HT1A receptor blocker WAY-100635 (below). **E)** Experimental paradigm for optogenetic stimulation during open field and TST. **F, G)** Optogenetic stimulation of 5-HT terminals in mOFC decreases the time in the outer zone and increases the time in the central zone of the open field in male ELS mice (mCherry: n = 4; ChR2: n = 6) (two-way ANOVA; outer zone, effect of group: F(1, 40) = 16.59, *p* = 0.0002, effect of time: F(4, 40) = 0.62, *p* = 0.65; central zone, effect of group: F(1, 40) = 16.59, *p* = 0.0002, effect of time: F(4, 40) = 0.59, *p* = 0.67). **H, I)** No effect of optogenetic stimulation in the time spent in outer or central zones of the open field in female ELS mice (mCherry: n = 4; ChR2: n = 6) (two-way ANOVA; outer zone, effect of group: F(1, 40) = 0.77, *p* = 0.39, effect of time: F(4, 40) = 0.47, *p* = 0.76). **J)** Optogenetic stimulation of 5-HT terminals in mOFC does not lead to a change in mobility of male ELS mice in TST (mCherry: n = 4; ChR2: n = 6) (two-way ANOVA; outer zone, effect of group: F(1, 16) = 0.58, *p* = 0.46, effect of time: F(1, 16) = 0.15, *p* = 0.71). **K)** Increased immobility in hChR2-mCherry expressing female ELS mice in TST during optogenetic stimulation of 5-HT terminals in mOFC (mCherry: n = 4; ChR2: n = 6) (two-way ANOVA; effect of group: F(1, 16) = 8.49, \**p* = 0.01, effect of time: F(1, 16) = 0.12, *p* = 0.73).

To determine whether DR^5-HT^ è mOFC pathway stimulation will affect stress coping strategy of male ELS mice and overcome the alterations in the active coping behavior observed in female ELS mice, we next performed TST in the absence or presence of blue light (470 nm, 20 Hz, 10 ms, 3 min epochs). While there were no significant differences in the mobility of male ELS mice in TST (two-way ANOVA; outer zone, effect of group: F(1, 16) = 0.58, *p* = 0.46, effect of time: F(1, 16) = 0.15, *p* = 0.71) (**Figure 6J**), female ELS mice expressing ChR2 had significantly increased immobility compared to those expressing mCherry only (two-way ANOVA; effect of group: F(1, 16) = 8.49, *p* = 0.01, effect of time: F(1, 16) = 0.12, *p* = 0.73) (**Figure 6K**). Although this may indicate a trend towards normalized mobility changes, the consequence of increased immobility with mOFC 5-HT terminal stimulation warrants further investigation.

Overall, stimulation of 5-HT neuron terminals in the mOFC significantly reduced the time spent in the outer zone of the open field and increased the time spent in the central zone, suggesting an anxiolytic effect in male ELS mice. While in females, the absence of an anxiolytic effect suggests that ELS-induced anxiety-like behavior may be influenced by different circuits.

## Discussion

Based on its significant brain-wide connectivity and role in circuit maturation, 5-HT plays a critical role in socioemotional regulation. Disruption of the development and activity of the 5-HT system during early life has persistent negative effects on affective behaviors. However, which neural circuits are involved in ELS-induced emotional dysregulation remains to be determined. Here, we used a modified LBN paradigm to recapitulate chronic stress during postnatal development in mice. We report that ELS induces altered maternal care, resulting in increased anxiety-like and impaired stress-related active coping behavior in female and male offspring during adulthood. Probing the ELS-induced network alterations revealed a significant disruption in the functional connectivity of the raphe nucleus in response to an aversive stimulus (footshock) predominantly in male ELS animals. The disruption of the raphe connectivity in ELS mice was accompanied by altered activity of the 5-HT neuron population to footshock. Further interrogation of two emotionally salient regions that receive dense 5-HT innervation revealed a significant increase in the activity of the OFC in response to footshock in ELS mice, without concomitant changes in the activity of CeA, which suggested that distinct 5-HT circuits may be differentially impaired in response to ELS. Fiber photometry in control and ELS mice revealed a significant decrease in the mOFC 5-HT release in avoidable- and unavoidable-threat environments. These impairments coincided with a disruption in the physiological 5-HT response of mOFC pyramidal neurons.

ELS has previously been associated with structural and functional alterations to fronto-limbic connectivity. Several key regions, including the hippocampus, medial prefrontal cortex and amygdala, have been particularly deemed susceptible to the effects of stress during the critical periods of development^58–60^. Given the strong significant connectivity of 5-HT neurons with regions and circuits implicated to be impaired in response to ELS, we sought to examine the functional connectivity of the raphe nucleus with 60 cortical and subcortical regions. Our data identified a shift towards anticorrelated functional connectivity of the raphe nucleus in response to ELS, with this effect being most prominent in males. This decrease in the correlation coefficient between ELS mice and controls was present across most regions paired with the raphe (42/59 in females, 56/59 in males). The OFC was among the most notable of these changes, with ELS decreasing the communication between this region and the raphe nucleus. This change is noteworthy as these two regions share reciprocal structural connections, through which the OFC contributes to the regulation of serotonergic inputs throughout the forebrain^61^. It is worth noting that the medial prefrontal cortex also shares this relationship with the raphe nucleus and changes in functional connectivity were also observed in this region. Further studies are needed to dissect the contributions of each of these pathways in behavioral changes observed with ELS.

It has recently been established that serotonin neuron population activity is influenced by environmental valence^41^ and encodes aversive states differentially^35, 38^. 5-HT neurons projecting to the OFC are preferentially inhibited by a negatively valanced stimulus such as footshock while those projecting to the CeA are activated by footshock^38^. Our data showing a selective disruption in the footstock-induced inhibitory response of 5-HT neurons suggested a subcircuit specific impairment in ELS mice in response to threat. In support of this, we observed an elevated footshock-induced activity of the OFC in ELS mice, indicating that this region is more sensitive to negative valence. Given the absence of significant activity differences in CeA, we characterized the alterations in 5-HT signaling induced by ELS in the OFC. The OFC is a highly conserved region involved in motivation, reward, and emotional regulation^62^, which are all disrupted in depressive disorders. Chronic exposure to early life adversities during infancy and to prenatal maternal depression results in reduced OFC cortical thickness observed many years later^63, 64^, in association with elevated depressive symptoms^65^. Our findings, which reveal disrupted 5-HT release during emotionally salient behaviors and impaired 5-HT physiology in the OFC of ELS mice, offer a mechanistic insight into the potential role of the OFC in mediating increased depressive symptoms following early life adversities.

ELS leads to long-lasting increased anxiety-like behavior in female and male mice, and altered coping strategies in female mice. One critical question that remains unanswered is whether impaired serotonin release and altered activity of the OFC in ELS mice contribute to the observed deficits in behavior. The role of the 5-HT signaling in the OFC was previously delineated in non-human primates and rodents in the context of reversal learning, cognitive flexibility and impulsivity and processing of emotional salience^61, 66–68^. While more studies are needed to elucidate how emotional behavior is modulated by 5-HT projections to the OFC, reducing 5-HT release in OFC of male mice has previously shown to increase anxiety-like behavior^38^. To our knowledge, ours is the first study that reports the effects of 5-HT neuron terminal stimulation in OFC in ELS mice. While stimulating the mOFC 5-HT terminals had an anxiolytic effect in male ELS mice, this approach was not sufficient to improve anxiety-like behavior of ELS females, necessitating further studies focused on the circuits underlying ELS-induced behavioral deficits in females. Moreover, exploring the impact of ELS-induced deficits in 5-HT signaling in the OFC on additional behaviors modulated by this region including impulsivity, cognitive flexibility, and reward processing could yield additional mechanistic insight into the broader effects of 5-HT dysfunction and its contribution to behavioral deficits associated with ELS.

In summary, our study presents a novel mechanistic understanding of the long-term impact of ELS-induced disruptions on brain circuits that are modulated by 5-HT inputs. Moreover, our findings reveal ELS-induced impairments in the DRN 5-HT è OFC pathway. While our network connectivity findings implicate additional pathways to be potentially impaired by ELS, the DRN 5-HT è OFC pathway represents a promising target for therapeutic intervention, given its critical role in reward and emotional regulation. Indeed, as the sex-dependent rescue of anxiety-like behavior by mOFC 5-HT terminal stimulation offers a potential therapeutic target for ELS-induced anxiety, our data also emphasize the importance of understanding the mechanisms of sex-dependent deficits underlying emotional dysregulation for the development of effective treatment strategies. Altogether, due to the increasing evidence of disrupted OFC connectivity in anxiety disorders^69–71^, the OFC has recently garnered a significant interest as a potential therapeutic target for novel neurostimulation treatments^72^. The top-down control mediated by this region onto other emotional circuits places it in a critical place and thus its activity to be maintained as normal is important. Therefore, our findings highlight the potential of combining targeted stimulation and pharmacotherapies to improve 5-HT neurotransmission as a promising approach for treating emotional dysregulation that arises from childhood stress.

## Acknowledgements

Funding for this study was provided by an CIHR Early Career Investigator Grant in Maternal, Reproductive, Child and Youth Health (#177446), a CFI-JELF fund (#42065) to DS and an Alberta Children’s Hospital Research Institute (ACHRI) Child Health and Wellness Grand Challenge Seedling Award to D.S. and J.R.E. RR received the Alberta Graduate Excellence Scholarship (AGES). K.A. received Eyes High Doctoral Recruitment Award. Y.R. was awarded the Harley Hotchkiss-Samuel Weiss Doctoral Scholarship. D.J.T. was awarded the NSERC Postgraduate Scholarship (PGS D). N.R. received Harley Hotchkiss-Samuel Weiss Postdoctoral Fellowship. Schematic images were created with BioRender.com.

## Author Contributions

D.S., R.R. and M.E. designed the experiments and wrote the manuscript. N.F.J. and M.T. performed the surgeries. R.R. and M.E. performed the behavioral, photometry and optogenetic experiments and performed the histological procedures. K.A., D.J.T and J.R.E. performed the network analysis. Y.R. contributed to the photometry and S.K. to the behavioral experiments. N.R. performed the electrophysiological experiments and analysis.

