## Supplemental Information for "Long-term impact of early life stress on serotonin connectivity"

**Supplemental Methods**

*Animals*

Female and male 10-20 week old mice on a C57BL6/J background were used for all experiments. Mice were housed in standard housing conditions on 12 h light/dark cycle (light on at 07:00) with water and food available *ad libitum*. All experimental procedures were performed in accordance with the guidelines established by the Canadian Council on Animal Care and were approved by the Life and Environmental Sciences Animal Care Committee at the University of Calgary. For behavioral characterization and functional connectivity experiments, Fos[2A-iCreER] (TRAP2) (JAX 030323) mice crossed with Ai14 reporter line (JAX 007914) (FosTRAP2::tdTomato mice) were used. For fiber photometry experiments to record $Ca^{2+}$ signal from 5-HT neurons, ePet-cre (JAX 012712) mice crossed with Ai148 reporter line (JAX 030328) (ePet1::Ai148 mice) were used. Wild-type C57BL6/J mice were used for fiber photometry experiments to record the iSeroSnFR signal.

*Limited Bedding and Nesting (LBN) Model*

On postnatal day (PND) 2, dams and pups were placed into custom designed cages fitted with a 3D printed plastic platform approximately 2 cm above the cage floor. The platform was perforated to allow urine and feces to pass through to the bedding underneath. Bedding sparingly covered the cage floor and half of a single cotton nestlet was provided as the nesting material. To prevent hypothermia, the temperature of the cage was maintained at $21^0$C from PND 2 to 4 using a thermostat and sub-cage heater. Control dams and pups were placed in similar cages on PND 2 but had access to regular bedding, nesting material and a see-through dome. The home-cage behavior of dams and pups in LBN and control cages were recorded during light and dark cycles with an upright camera mounted to the lid of each cage. Both control and LBN dams and pups were returned to a new cage with a regular bedding and nesting material on PND 12.

*Behavioral Analysis*

All behavioral tests were performed between 9 a.m. and 3 p.m. Mice were acclimatized to the testing environment for 30 min prior with a white noise generator operating in the background for the entire duration of the experiments. Each apparatus was cleaned with 70% ethanol and wiped down with a paper towel before testing each animal. All test sessions were recorded via ANY-Maze video tracking and behavioral analysis software (Stoelting Co.).

<u>Open Field Test (OFT)</u>: OFT was performed to assess anxiety-like behavior. Mice were placed in an acrylic 40 cm X 40 cm apparatus and were allowed to explore for 15 min. Videos were recorded and the time spent in each zone, and distance traveled were analyzed using ANY-maze (Stoelting Co.).

<u>3-Chamber Social Interaction Test:</u> The 3-chamber test was performed to assess sociability. Mice were placed in an apparatus (105 cm X 45 cm) with two plexiglass dividers, splitting the chamber into three equal zones. During habituation, each mouse was placed in the middle zone with dividers in place for 5 min. After dividers were removed, mice were allowed to explore the apparatus freely for 5 min. During test stage, an age-, strain- and sex- matched novel mouse was placed on one side of the chamber (social chamber) confined in an inverted pencil cup while the other chamber contained an empty cup (non-social chamber). Each mouse was first placed in the middle zone for 5 min followed by removal of the dividers and allowing mice to explore for 10 min. Videos were recorded and the time spent in social and non-social interaction zones (a circular zone with 2 cm radial distance to the pencil cup) was analyzed using ANY-maze (Stoelting Co.). Sociability index was calculated as (time in social interaction zone – time in non-social interaction zone)/ (time in social interaction zone + time in non-social interaction zone).

<u>Tail Suspension Test (TST)</u>: TST was performed to assess active and passive stress coping strategies. 17 cm tape fragments were used to suspend each mouse from its tail from a metal rod hanging 60 cm above the surface for a duration of 6 min. Tails were gently placed in a 3 cm long plastic cylinder (1 cm in diameter, 2 gr in weight) to prevent tail climbing behavior. Videos were

recorded using ANY-maze (Stoelting Co.). Mobility was scored manually by an experimenter blinded to the conditions.

*Functional Connectivity of the Dorsal Raphe*

The brain-wide functional connectivity of the dorsal raphe nucleus was assessed in response to acute stress using IEG-based analyses of regional coactivation. Acute stress was induced with a protocol in which mice received a series of 10 foot-shocks (0.5 mA) with an interval of 30-seconds between shocks. Following the conclusion of the acute stress protocol, mice were injected with a dose of 0.1 mg/g 4-hydroxytampoxifen (HelloBio, HB6040-50mg) and left undisturbed in their home cages for 3 days to allow for labelling of cells which were active during the stress protocol. Mice were transcardially perfused with 0.1 M phosphate buffered saline (PBS) followed by 4% formaldehyde. Brains were extracted and incubated in 4% formaldehyde for 24 h before being cryoprotected in a 30% sucrose solution until no longer buoyant (48 – 72 h). Cryoprotected brains were then sectioned 40 μm thick on a cryostat (Leica) in a 6 series and stored at -20 °C in a buffered antifreeze solution containing 30% ethylene glycol and 20% glycerol in 0.1 M PBS.

Excess antifreeze solution was washed from the tissue (3 X 10 min in 0.1 M PBS) and sections were counterstained with DAPI. Tissue was mounted to glass slides and coverslipped with PVA-DABCO mounting medium. An OLYMPUS VS120-L100-W slide scanning microscope was used to collected images of the brain-wide expression of the c-fos – dTomato label and the DAPI counterstain. Fluorescently labelled c-fos positive cells were segmented based on fluorescent intensity and morphological characteristics using user-guided machine-learning-based *Ilastik* software[1]. Images of the segmented c-fos positive cells and corresponding DAPI counterstains were reconstructed into 3D volumes for each mouse using TissueMaker (MBF Bioscience). These volumed were then aligned to the Allen Brain Institute's mouse brain reference atlas using NeuroInfo (MBF Bioscience) and the density of c-fos positive cells was determined across 60 brain regions.

Functional connectivity analyses were performed using MATLAB analyses derived from previously described analyses of IEG-based functional connectivity[2,3]. MATLAB scripts were modified to accommodate data outputs from NeuroInfo. Within each group, the expression density of c-fos positive cells was cross correlated between all possible pairs of regions using a Pearson's correlation to generate functional connectivity matrices. The column of the functional connectivity matrix detailing the functional connectivity of the dorsal raphe was isolated in each group and the number of correlations with a negative Pearson's *R* correlation coefficient (anticorrelations) and the distribution of Pearson's *R* correlation coefficients was compared across groups.

*Stereotaxic surgeries*

Mice were anaesthetized using isoflurane (5% during induction, 2% for maintenance) and oxygen (0.4 L/min). Eye gel ointment was applied for lubricating the eyes during surgery. Mice received meloxicam (5 mg/kg, s.c.) for analgesia and saline (s.c.) for fluid support. Skin above the head was shaved and cleaned with 70% ethanol and iodine. In ePet1::Ai148 mice, a small hole in the skull was opened using an electric drill and monofiber optic cannulae (4 mm length, 400 um diameter, 0.46 NA; Neurophotometrics) were implanted at an angle of 30˚ above the DRN according to the following stereotaxic coordinates (in mm): AP: -6.27, ML: 0, DV: -4.04 from Bregma. For 5-HT sensor experiments, wild-type C57BL6/J mice received a unilateral injection of 400 nl AAV2/9-CAG-iSeroSnFR-NLG (1.0 x E12 vg/ml; Canadian Neurophotonics Platform Viral Core Facility; RRID:SCR_016477) into the mOFC (AP: 2.34, ML: 0.3, DV: 2.5 mm from Bregma) using a glass micropipette attached to a Nanoject III infusion system (Drummond Scientific). Monofiber optic cannulae (4 mm length, 400 um diameter, 0.46 NA; Neurophotometrics) were implanted above the mOFC (AP: 2.34, ML: 0.3, DV: 2.4 mm from Bregma). For optogenetic experiments, the DRN of ePet1::Cre mice was infused with 800 nl AAV2/9-EF1a-DIO-mCherry or (6.0 x E12 vg/ml; Canadian Neurophotonics Platform Viral Core Facility; RRID:SCR_016477). Optic cannulae (3 mm length, 200 um diameter, 0.37 NA; Neurophotometrics) were implanted bilaterally at a 21˚ angle above the mOFC (AP: 2.34, ML: ±1.25, DV: -2.6 from Bregma). The cannulae were secured by an initial application of superglue followed by layer of dental cement. Behavioral experiments were performed at least 3 weeks after surgery.

*In vivo $Ca^{2+}$ imaging of 5-HT neurons*

The activity of 5-HT neuron population was recorded in ePet1::Ai148 mice during exposure to ten footshocks (0.5 mA, 2 sec each, 30 sec apart). A Doric Lenses fiber photometry system consisted of two Light Emitting Diodes (LEDs) including the 465 nm wavelength for GCaMP6f stimulation and 405 nm wavelength as the isosbestic control signal. Light was coupled to a low fluorescence patch cord (1.5 m, 0.48 NA, Doric Lenses). 465 nm LED was modulated at 211 Hz, passed through a GFP emission filter and focused onto a photodetector (Model 2151, Newport). Prior to any experiment, a photodetector (THORLabs) was used to measure the light intensity (μW) for both wavelengths to ensure consistency of light delivery (30 μW) for all experiments. Behavior and photometry signals were synchronized using the “time-to-live” (TTL) trigger function on the fiber photometry console that operated simultaneously with ANY-maze behavioral tracking software. Data were extracted and analyzed using custom-written script in MATLAB (Mathworks) as previously described[4]. Accordingly, isosbestic 405 nm data was fit to a biexponential decay which was linearly scaled to the raw 465 nm data. A Fourier ΔF/F baseline correction of the data was performed to remove artifacts related to photobleaching or motion. Z-scored ΔF/F traces were calculated and analyzed.

*In vivo measurement of 5-HT dynamics*

4 weeks after surgery, fiber photometry was performed to measure serotonin release during open field and tail suspension tests. Prior to behavioral tasks, mice were habituated to being connected to the fiber optic patch cord for 3 consecutive days. The *in vivo* photometry recordings for open field and tail suspension test were conducted using a Neurophotometrics FP3002 system controlled by Bonsai software. The Neurophotometrics system simultaneously received TTL pulses generated by ANY-maze behavioral tracking software during the behavior task. A 470 nm LED delivered an excitation wavelength of light to collect 5-HT sensor-dependent data. 5-HT

sensor-independent isosbestic signals were simultaneously captured by excitation with 415 nm LED. The region calibration for the patch cord and the power calibration (μW) for both wavelengths to ensure consistency of 100 μW light delivery were performed before the experiment. All wavelengths were interleaved and collected using Bonsai with 30 Hz sampling rate (30 frames per second). 415 nm data was fit to a biexponential decay to correct for photobleaching and the resulting vector was used to linearly scale 470 nm data. To calculate ΔF, these linearly scaled 5-HT sensor-dependent data were subtracted from the raw unprocessed calcium-dependent data. The resulting values were divided by the linearly scaled calcium-dependent data to generate a ΔF/F trace. ΔF/F during specific behaviors and at different behavioral epochs were identified using custom-written script in MATLAB[5]. ΔF/F traces corresponding to -2 s to 3 s entry period while each mouse transitioned from the outer to the intermediate zone of the open field and -2 s to 2 s period when mice transitioned from mobility to immobility within the last 4 min of the TST were calculated. The resulting ΔF/F traces were also analyzed for area under the curve (AUC), and the averaged z-score. AUC was operationally defined as the summed area between the X-axis and the ΔF/F trace.

*In vivo optogenetic stimulation*

Laser driver (Opto Engine LLC) was connected to a 1 x 2 fiber optic rotary joint (Doric Lenses) attached to a dual fiber-optic patch cord. Trains of blue light (470 nm, 10 mW) at 20 Hz, 10 ms were used every 3 min throughout the open field (15 min) and tail suspension (6 min) tests, starting with a 3 min light off epoch. Light was delivered via ANY-Maze software-controlled AMi-2 optogenetic interface (Stoelting Co.) connected to the laser driver.

*Immunohistochemistry*

For immunostaining, sections were washed 3 X 10 min with 0.1 M PBS and incubated for 24 h with rabbit anti-c-fos (1:1000, SYSY), mouse anti-mCherry (1:500, BioLegend) or goat anti-5-HT (1:500, Abcam) in 0.1 M PBS containing 3% Normal Donkey Serum (NDS) and 0.3% Triton-X-100. The next day, sections were washed 3 X 10 min with 0.1 M PBS and incubated with donkey anti-rabbit Alexa Fluor 647 (1:500, Invitrogen), anti-mouse Alexa Fluor 647 (1:500, Jackson ImmunoResearch), anti-goat Alexa Fluor 488 (1:500, Jackson ImmunoResearch) or anti-goat Alexa Fluor 594 (1:500, Invitrogen) in dark for 24 h. Following secondary antibody incubation, sections were washed 3 X 10 min with 0.1M PBS and were incubated with DAPI (1:500, HelloBio) in PBS for 5 min. After 2 X 10 min washes with 0.1 M PBS, sections were mounted on glass slides and coverslipped with PVA-DABCO mounting medium. For virus and optical fiber validation, sections were imaged using an Olympus FV3000 confocal microscope.

*Quantification of c-Fos density using FASTMAP*

Sections were scanned at 10X magnification using a slide scanner (OLYMPUS VS120-L100-W; Richmond Hill, ON, CA). From these images, c-fos positive cells were segmented from background based on fluorescent intensity and label morphology using *Ilastik*[1]. Images were then registered to the Allen Brain Institute's mouse brain reference atlas to assess the expression density of segmented c-fos positive cells in the OFC and CeA regions using FASTMAP[6]. The expression density of these cells has been presented as the number of c-fos positive cells divided by the area of the region ($um^2$) was analyzed for each brain.

*Ex vivo electrophysiology*

400 um slices comprising the mOFC were obtained from control and ELS mice using a vibratome and recovered in aCSF (128 mM NaCl, 10 mM D-glucose, 26 mM NaHCO3, 2 mM CaCl2, 2 mM 589 MgSO4, 3 mM KCl, 1.25 mM NaH2PO4, pH 7.4) with 95% $O_2$/5% $CO_2$ at 30$^o$C. Recordings were performed in oxygenated aCSF flowing at a rate of 3-4 ml/min. The internal patch solution contained 120 mM potassium gluconate, 10 mM HEPES, 5 mM KCl, 2 mM MgCl2, 4 mM K2-ATP, 0.4 mM Na2-GTP, 10 mM Na2-phosphocreatine, with pH adjusted to 7.3. mOFC pyramidal neurons were visualized using IR-DIC using an Olympus BX51WI microscope and identified based on their characteristic pyramidal cell body. Whole-cell recordings were obtained in voltage clamp and current clamp mode using a MultiClamp 700B amplifier (Molecular Devices). 5-HT (10 uM, Sigma) was bath applied for 30 sec to record current response while cells were held at -75

mV in voltage-clamp mode. For *ex vivo* optogenetic experiments, current was injected to stimulate action potentials at 1-2 Hz in current-clamp mode. Light (470 nm, 20 Hz, 10 ms) was turned on for 5 sec to stimulate 5-HT release. 30 nM WAY-100635 (Sigma) was applied for 5 min to block 5-HT1A receptor signaling. Data were filtered at 4 kHz and digitized at 20 kHz using Digidata 1550B and Clampex software (Molecular Devices) and analyzed using Clampfit software.

*Statistical analysis*

Statistical analysis was performed using GraphPad Prism 9. Data were analyzed using two-way ANOVA or unpaired t-test. Following significant interactions in two-way ANOVA , Tukey's multiple comparisons tests were used. Statistical significance was set to $p < .05$. All data and figures were presented as mean ± standard error of the mean (SEM). Statistical analyses for the generation of functional connectivity matrices were performed using MATLAB (R2020a Update 1). Matrices were generated using Pearson's correlation. Assessment of the distribution of Pearson's *R* correlation coefficient was performed using the Kolmogorov-Smirnov test in GraphPad Prism 9. Circle plots were generated using the circular Graph MATLAB function. Some of the schematics were generated using images from BioRender.

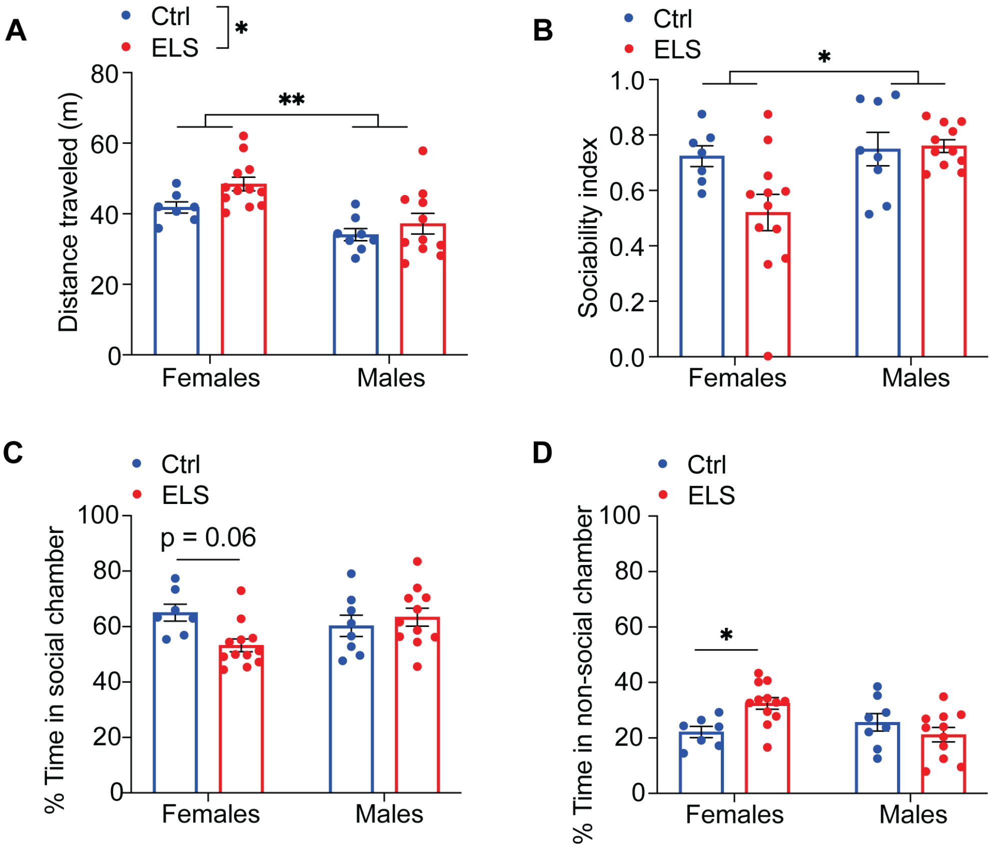


**Figure S1. Locomotor activity and sociability in control and ELS mice during adulthood. A)** Control and ELS female and male mice traveled comparable distance in the open field test, with females showing an overall greater locomotor activity, compared with males (two-way ANOVA; effect of group: $F(1, 34) = 4.31$, $p = 0.04$, effect of sex: $F(1, 34) = 16.4$, $p < 0.001$). **B)** Males showed greater sociability than females, with no significant difference between control and ELS groups (two-way ANOVA; effect of group: $F(1, 34) = 3.21$, $p = 0.08$, effect of sex: $F(1, 34) = 6.11$, $p = 0.019$). **C)** Female ELS mice showed a tendency for spending less time in the social chamber (two-way ANOVA; group X sex interaction: $F(1, 34) = 5.62$, $p = 0.024$, Tukey's multiple comparisons test: Female ctrl vs ELS $p = 0.067$; female ELS vs male ELS $p = 0.06$, all other *p*'s $> 0.1$). **D)** Female mice spent significantly greater time in the non-social chamber, compared to controls (two-way ANOVA; group X sex interaction: $F(1, 34) = 8.18$, $p = 0.007$, Tukey's multiple comparisons test: Female ctrl vs ELS $p = 0.04$; female ELS vs male ELS $p = 0.007$, all other *p*'s $> 0.1$).

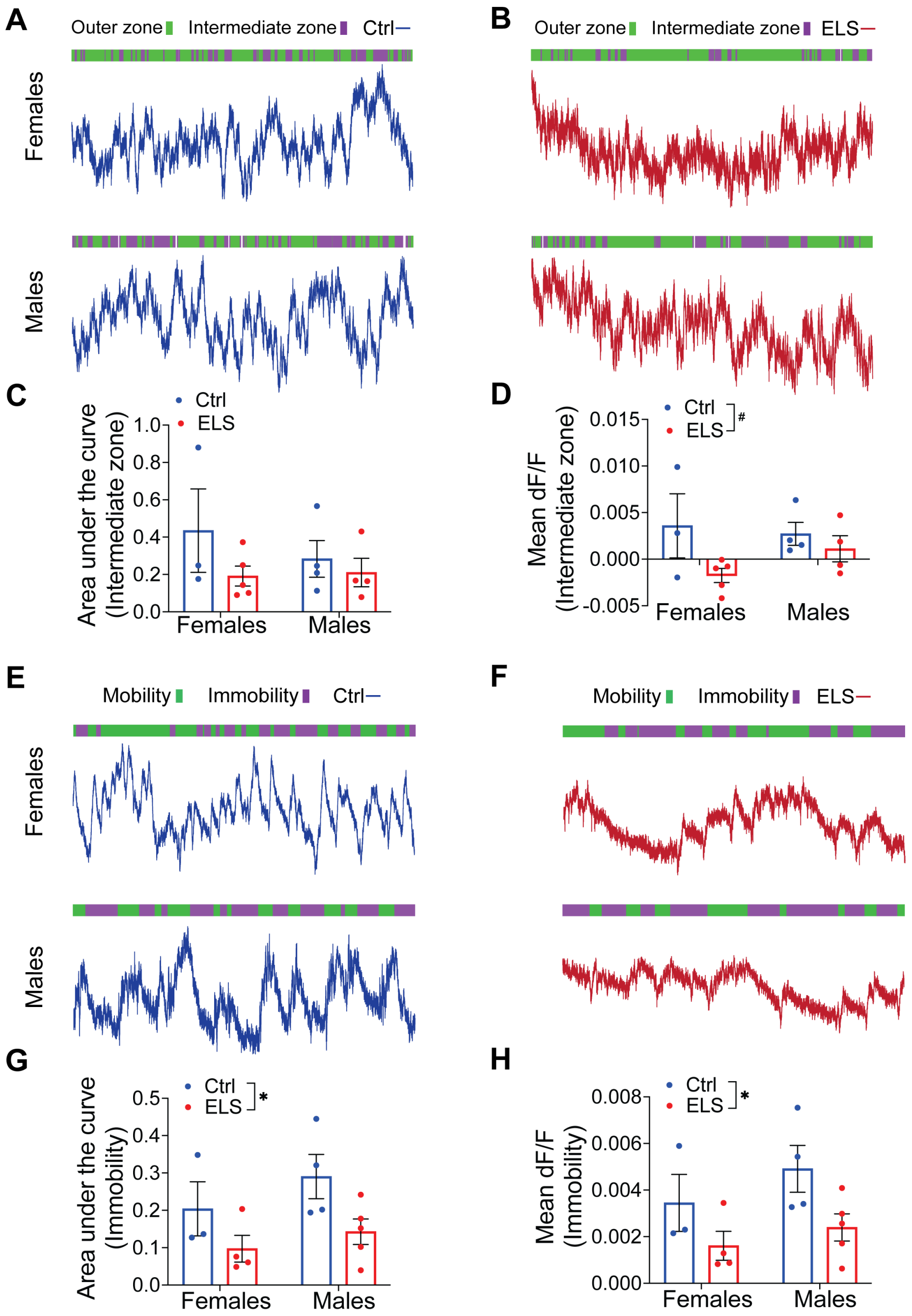


**Figure S2. 5-HT release in mOFC during open field and TST. A, B)** Representative photometry traces of the iSeroSnFR signal in the mOFC of female and male control and ELS mice during open field test are shown. Behavior bouts corresponding to the position of mice in different zones time locked to the photometry signal are shown above. **C)** The area under the curve of the iSeroSnFR signal of male and female control and ELS mice in the intermediate zone of the open field (two-way ANOVA; effect of group: $F(1, 12) = 2.14$, $p = 0.17$; effect of sex: $F(1, 12) = 0.38$, $p = 0.55$). **D)** The mean amplitude of the iSeroSnFR signal of male and female control and ELS mice in the intermediate zone of the open field (two-way ANOVA; effect of group: $F(1, 12) = 4.49$, $^{\#}p = 0.05$; effect of sex: $F(1, 12) = 0.38$, $p = 0.55$). **E, F)** Representative photometry traces of the iSeroSnFR signal in the mOFC of female and male control and ELS mice during TST are shown. Behavior bouts corresponding to the mobility and immobility are shown above. **G)** The area under the curve of the iSeroSnFR signal of male and female control and ELS mice during immobility in the TST (two-way ANOVA; effect of group: $F(1, 12) = 6.7$, *$p = 0.02$; effect of sex: $F(1, 12) = 1.8$, $p = 0.2$). **H)** The mean amplitude of the iSeroSnFR signal of male and female control and ELS mice during immobility in the TST (two-way ANOVA; effect of group: $F(1, 12) = 6.8$, $^{*}p = 0.02$; effect of sex: $F(1, 12) = 1.81$, $p = 0.2$).

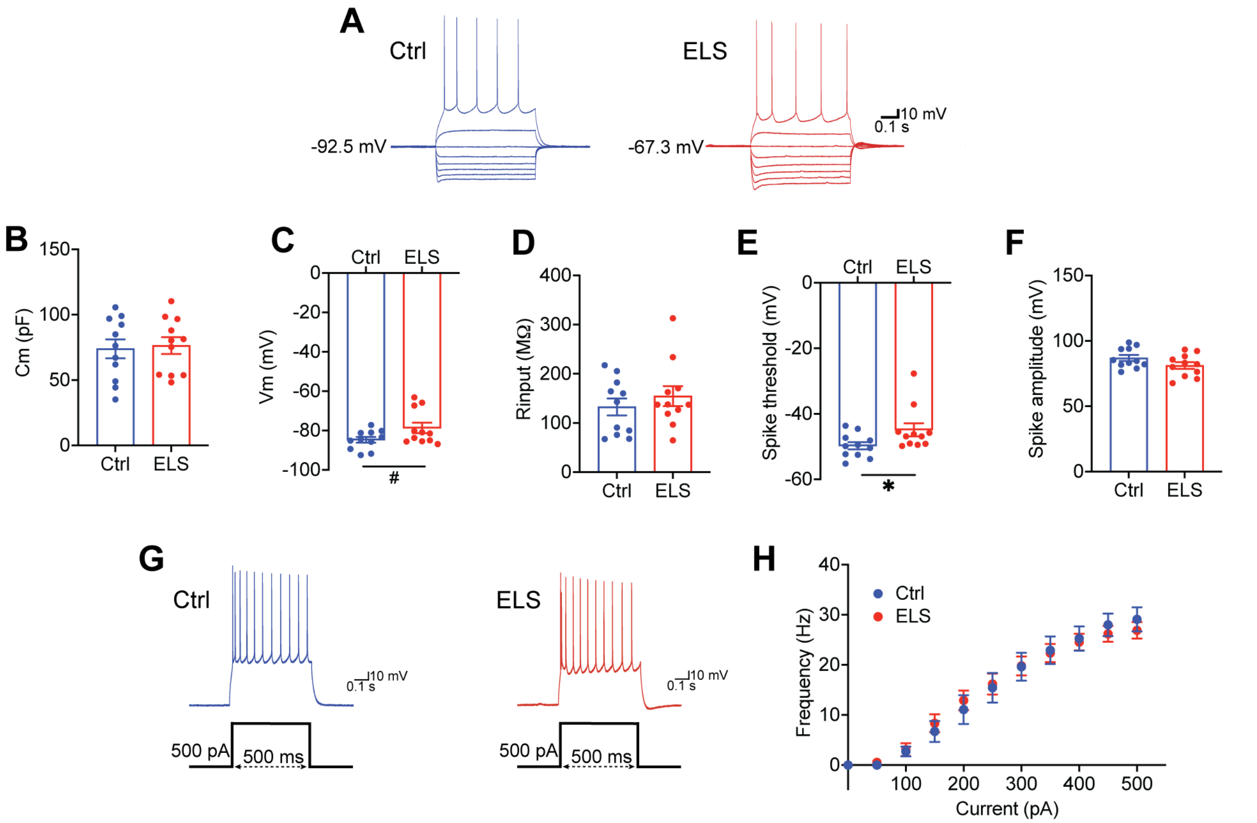


**Figure S3. Passive membrane characteristics and excitability of mOFC pyramidal neurons of control and ELS mice. A)** Representative current clamp recordings from mOFC pyramidal neurons in response to depolarizing and hyperpolarizing current steps from a control and an ELS mouse. **B)** Membrane capacitance (Cm; pF) of mOFC pyramidal neurons of control and ELS mice (unpaired t-test, $p = 0.8$) (4 control male mice, n = 11 neurons; 3 ELS male mice, n = 11 neurons). **C)** Membrane voltage (Vm; mV) of mOFC pyramidal neurons of control and ELS mice (unpaired t-test, $^{\#}p = 0.06$). **D)** Input resistance (Rinput; MΩ) of mOFC pyramidal neurons of control and ELS mice (unpaired t-test, $p = 0.8$). **E)** The spike threshold (mV) of mOFC pyramidal neurons of ELS mice is significantly reduced compared to those of controls (unpaired t-test, $^{*}p = 0.046$). **G)** Representative current clamp recordings of mOFC pyramidal neurons from a control and an ELS mouse in response to 500 pA depolarizing current input. **H)** Input-output curve showing spike frequency (Hz) of mOFC pyramidal neurons in response to a series of depolarizing current injections. The frequency of action potentials is comparable between control and ELS mice (two-way ANOVA; effect of group: $F(1, 220) = 0.0$, $p > 0.99$; effect of current: $F(10, 220) = 60.91$, $p < 0.01$).

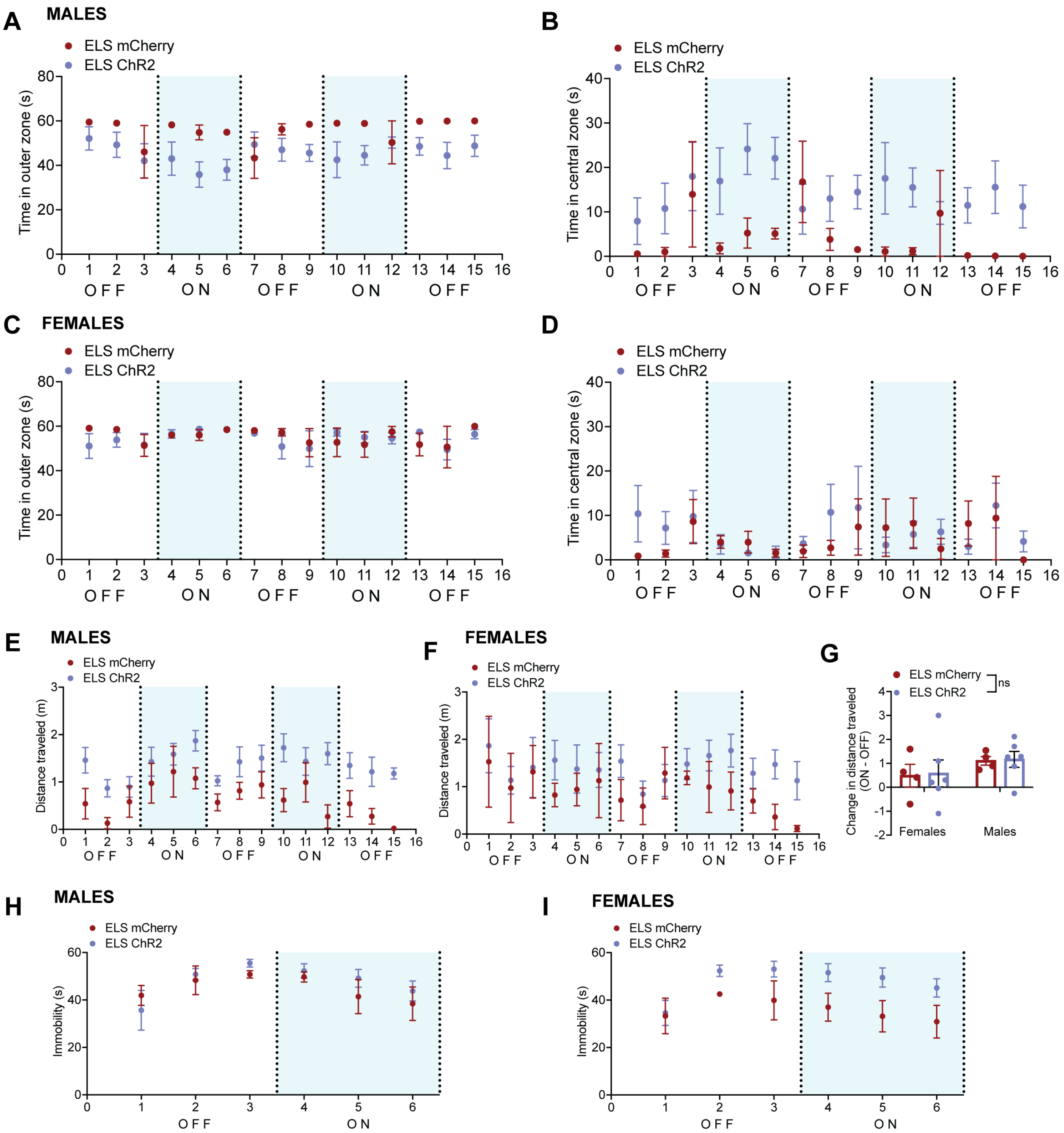


**Figure S4. Optogenetic inhibition of 5-HT terminals in the mOFC during open field and TST. A, B)** The time spent by mCherry- and ChR2-expressing ELS male mice in the outer and central zones of the open field during light off and on epochs over time are shown. **C, D)** The time spent by mCherry- and ChR2-expressing ELS female mice in the outer and central zones of the open field during light off and on epochs over time are shown. The amount of distance traveled in the open field by mCherry- and ChR2-expressing ELS male **(E)** and female **(F)** mice during light off and on epochs over time are shown. **G)** The effect of optogenetic stimulation on distance traveled in female and male mice (two-way ANOVA; effect of group: $F(1, 16) = 0.03$, $p = 0.87$; effect of sex: $F(1, 16) = 1.75$, $p = 0.2$). The time spent immobile in the TST by mCherry- and ChR2-expressing ELS male **(H)** and female **(I)** mice during light off and on epochs over time are shown.

Supplemental Table 1

| ABBREVIATION | FULL NAME |
|---|---|
| MO1 | Primary motor area |
| MO2 | Secondary motor area |
| FRP | Frontal pole |
| ACA | Anterior cingulate area |
| PL | Prelimbic area |
| IL | Infralimbic area |
| ORB | Orbital area |
| AI | Agranular insular area |
| AON | Anterior olfactory nucleus |
| TT | Taenia tecta |
| DP | Dorsal peduncular area |
| PIR | Piriform area |
| EP | Endopiriform nucleus |
| STR | Striatum |
| SSp | Primary somatosensory area |
| GUS | Gustatory areas |
| CLA | Claustrum |
| PAL | Pallidum |
| SSs | Supplemental somatosensory area |
| VISC | Visceral area |
| HY | Hypothalamus |
| COA | Cortical amygdalar area |
| PAA | Piriform-amygdalar area |
| TH | Thalamus |
| AHN | Anterior hypothalamic nucleus |
| NLOT | Nucleus of the lateral olfactory tract |
| BMA | Basomedial amygdalar nucleus |
| ATN | Anterior group of the dorsal thalamus |
| MED | Medial group of the dorsal thalamus |
| BLA | Basolateral amygdalar nucleus |
| RSC | Retrosplenial area |
| CA3 | Field CA3 |
| DG | Dentate gyrus |
| CA1 | Field CA1 |
| CA2 | Field CA2 |
| LA | Lateral amygdalar nucleus |
| PVH | Paraventricular hypothalamic nucleus |
| VMH | Ventromedial hypothalamic nucleus |
| PTLp | Posterior parietal association areas |
| LAT | Lateral group of the dorsal thalamus |
| AUD | Auditory areas |
| TEa | Temporal association areas |

| | |
|---|---|
| PERI | Perirhinal area |
| ECT | Ectorhinal area |
| ENT | Entorhinal area |
| STN | Subthalamic nucleus |
| VIS | Visual areas |
| SUB | Subiculum |
| PA | Posterior amygdalar nucleus |
| PAG | Periaqueductal gray |
| TR | Postpiriform transition area |
| ProS | Prosubiculum |
| SNr | Substantia nigra |
| VTA | Ventral tegmental area |
| POST | Postsubiculum |
| SC | Superior colliculus |
| RAPHE | Raphe nuclei |
| CB | Cerebellum |
| PRE | Presubiculum |
| IC | Inferior colliculus |
